# Oligodendrocyte secreted factors shape hippocampal GABAergic neuron transcriptome and physiology

**DOI:** 10.1101/2020.11.09.374306

**Authors:** Elisa Mazuir, Louis Richevaux, Merie Nassar, Noémie Robil, Pierre de la Grange, Catherine Lubetzki, Desdemona Fricker, Nathalie Sol-Foulon

## Abstract

Oligodendrocytes form myelin for central nervous system axons and release factors which signal to neurons during myelination. Here, we ask how oligodendroglial factors influence hippocampal GABAergic neuron physiology. In mixed hippocampal cultures GABAergic neurons fired action potentials of short duration and received high frequencies of excitatory synaptic events. In purified neuronal cultures without glial cells, GABAergic neuron excitability increased and the frequency of synaptic events decreased. These effects were largely reversed by adding oligodendrocyte conditioned medium. We compared the transcriptomic signature with the electrophysiological phenotype of single neurons in these three culture conditions. Genes expressed by single pyramidal or GABAergic neurons largely conformed to expected cell-type specific patterns. Multiple genes of GABAergic neurons were significantly downregulated by the transition from mixed cultures containing glial cells to purified neuronal cultures. Levels of these genes were restored by the addition of oligodendrocyte conditioned medium to purified cultures. Clustering genes with similar changes in expression between different culture conditions revealed processes affected by oligodendroglial factors. Enriched genes are linked to roles in synapse assembly, action potential generation and transmembrane ion transport, including of zinc. These results provide new insight into the molecular targets by which oligodendrocytes influence neuron excitability and synaptic function.

---

Communication between oligodendrocytes and neurons is crucial for circuit maturation but still not completely understood. The fast transmission of action potentials relies on insulating properties of myelin sheath which is interrupted at nodes of Ranvier, small axonal domains highly enriched in voltage-gated Na^+^ channels (Sherman and Brophy 2005). The profile of myelination and nodes of Ranvier controls the timing of impulse transmission, critical for coincident arrival of synaptic inputs transmitted by multiple axons in sensory systems (Seidl 2014; Freeman et al. 2016; Arancibia-Cárcamo et al. 2017; Monje 2018). Both oligodendrocytes and oligodendrocyte precursor cells (OPCs or NG2 cells) sense neuronal activity, which triggers their differentiation and maturation into myelinating oligodendrocytes (Barres & Raff, 1993; Demerens et al. 1996). Adaptive myelination acts to reinforce selected circuits during learning (McKenzie et al. 2014; Bechler et al. 2018; Monje 2018; Stedehouder et al. 2018). Oligodendrocytes also release lactate to provide metabolic support to axons (Fünfschilling et al. 2012; Lee et al. 2012; Saab et al. 2016).

Factors secreted by oligodendrocytes induce early formation of node-like clusters, termed prenodes, enriched in Na_v_ channels, Nfasc186 and Ankyrin-G, along the axons of retinal ganglion cells and some hippocampal GABAergic neurons (parvalbumin or somatostatin immunopositive) before myelination (Kaplan et al. 1997; Freeman et al. 2015; Bonetto et al. 2019; Dubessy et al. 2019). These early clusters are associated with an increased axonal conduction velocity along GABAergic axons (Freeman et al. 2015). In addition, factors secreted by oligodendrocytes or their precursor cells close to the soma of pyramidal neurons modulate glutamatergic neurotransmission and restrain high-frequency firing (Sakry et al. 2014; Birey et al. 2015; Battefeld et al. 2016; Jang et al. 2019; Xin et al. 2019). Moreover, oligodendroglial exosomes and OPC secreted protein NG2 mediate glia signaling to neurons (Frühbeis et al. 2013; Sakry et al. 2015). The identity of molecular targets by which oligodendrocyte secreted factors affect GABAergic neuron excitability and synaptic interactions remain unknown.

The present study aimed to identify targets of oligodendrocyte mediated regulation of GABAergic hippocampal neurons. Electrophysiological phenotypes of rat hippocampal neurons were compared in mixed cultures, (with glial cells, CTRL) and purified neuron cultures (without glial cells, PUR). We then tested the effects of adding oligodendrocyte conditioned medium (OCM) to purified cultures. OCM tended to reverse changes in GABAergic neuron physiology and anatomy induced by eliminating glial cells from cultures Single-cell RNA-sequencing of GABAergic neuron cytoplasm collected in patch electrodes let us explore molecular targets of OCM-induced regulation. RNA-seq analysis was validated by the presence of cell-type specific genes, including those for subclasses of GABAergic neurons. Major targets of oligodendrocyte factor signaling to GABAergic neurons included ion channels and transporters contributing to the regulation of membrane potential and action potential generation as well transmembrane transport of zinc.

## MATERIALS AND METHODS

### Animals

Care and use of rats in all experiments conformed to institutional policies and guidelines (UPMC, INSERM, and European Community Council Directive 86/609/EEC). The following rat strains were studied: Sprague-Dawley or Wistar rats (Janvier Breeding Center) and VGAT-Venus Wistar rats in which a green fluorescent protein variant is selectively expressed in GABAergic cells (Uemastu et al. 2008). We assume that fluorescent cells in cultures correspond to GABAergic neurons.

### Culture Media

We used the following culture media. NM, neurobasal medium (21103ium (2Gibco) supplemented with 0.5 mM L-glutamine, B27 (1×; Invitrogen), and penicillin-streptomycin (100 IU/mL). BS, Bottenstein-Sato medium: DMEM Glutamax supplemented with transferrin (50 μg/mL), albumin (50 μg/mL), insulin (5 μg/mL), progesterone (20 nM), putrescine (16 μg/mL), sodium selenite (5 ng/mL), T3 (40 ng/mL), T4 (40 ng/mL) and PDGF (10 ng/ml).

### Preparation of oligodendrocyte conditioned medium

Glial cell cultures were prepared from cerebral cortices of P2 Wistar rats as described previously (Mazuir et al. 2020). After meninges were removed, cortices were incubated for 35 min in papain (30 U/mL; Worthington), supplemented with L-cysteine (0.24 mg/mL) and DNase (50 μg/mL) in DMEM at 37°. They were then mechanically homogenized and passed through a 0.70 μm filter. Cells were re-suspended in DMEM with 10% FCS and 1% penicillin-streptomycin. After 7 to 14 days *in vitro* (DIV), oligodendroglial lineage cells were purified from glial cell cultures which initially contain astrocytes and microglial cells. After cultures were shaken overnight at 230 rpm and 37°C, overlying oligodendroglial and microglial cells could be selectively detached. Microglia were then eliminated by differential adhesion (McCarthy and de Vellis 1980). Collected cells were incubated in dishes for 15 min. Non-adherent cells were retrieved and centrifuged in DMEM for 5 min at 1500 rpm. They were re-suspended and seeded at a density of 1.5×10^5^/cm^2^ on Polyethylene-imine (PEI)-coated dishes with BS medium and 0.5% PDGF. Immunostaining showed that 90 ± 4% of cells were positive for the oligodendroglial marker O4^+^, 7.2 ± 2.5% were GFAP^+^ astrocytes and 4.6 ± 0.7% were CD11b^+^ immuno-positive microglial cells (Mazuir et al. 2020). Medium from these cultures was collected after 48 hours, filtered (0.22 μm) and stored for use as oligodendrocyte conditioned medium.

### Neuronal cultures

Experiments were performed in three different culture conditions (Fig. 1A). Control (CTRL) was mixed hippocampal cultures containing neurons, astrocytes and oligodendrocytes. They were prepared from E18 rat embryos and have been characterized (Freeman et al. 2015). Purified neuron cultures (PUR) were prepared by adding anti-mitotic agents (FdU and U 5μM) for 12 hours, starting at 24 hours after dissection. Immunostaining showed these cultures contained less than 5% astrocytes and virtually no oligodendrocytes. In OCM cultures oligodendrocyte conditioned medium was added to purified neuron cultures. Conditioned medium (500 μl/well) was added at 3 days *in vitro* (DIV). One-third of the medium was replaced with neurobasal medium (NM) at 7 DIV, and then twice a week. Axon initial segments were visualized by 20 min exposure to an anti-Nfasc antibody (clone A12/18, Antibodies Incorporated) coupled to Alexa 594 (using Apex antibody labeling kit, ref A10474, Thermofisher) before recordings.

**Figure 1:**
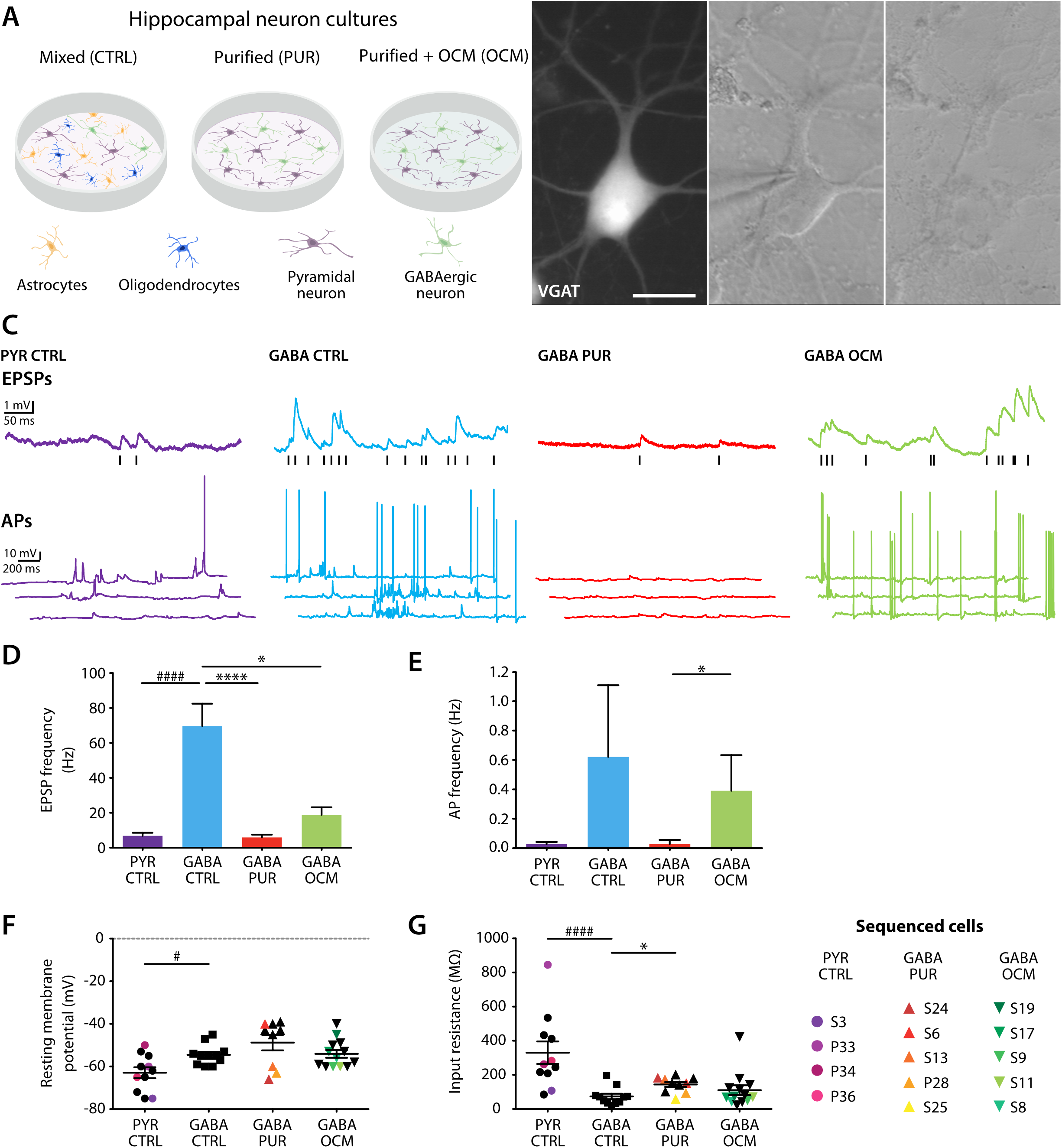
Patch clamp recording and cytosol harvesting of hippocampal neurons. (A) Schematic representation of the culture conditions: neurons from mixed cultures (CTRL), containing hippocampal neurons and glial cells, were compared with those of purified cultures (PUR), which in some cases were treated with oligodendroglia conditioned medium (OCM). (B) Soma and proximal dendrites of a green fluorescent GABAergic neuron (VGAT+ cell; left). Patch pipette sealed to the neuronal membrane for electrical recording (middle). After aspiration of the cytosolic content (right). Scale bar: 20 μm (C) Representative voltage recordings of pyramidal and GABAergic neurons at 17 DIV in different culture conditions. Top, excitatory postsynaptic potentials (EPSPs), indicated by black lines. Bottom, spontaneous action potential (AP) firing. (D, E) EPSP (D) and AP (E) frequencies measured from neurons in different conditions. EPSP, PYR CTRL vs. GABA CTRL, p<0.0001 (Mann-Whitney test #); GABA CTRL vs. GABA PUR, p < 0.0001; GABA CTRL vs. GABA OCM, p=0.0123. AP, GABA PUR vs. GABA OCM, p=0.0126 (Kruskal-Wallis and Dunn’s *post hoc \**). (F, G) Resting membrane potential (F) and input resistance (G) of recorded neurons in different conditions. Color symbols show cells from which sequence data was derived. Mann-Whitney test for PYR CTRL vs. GABA CTRL significance levels indicated with #, Kruskal-Wallis and Dunn’s *post hoc* for GABA CTRL vs. PUR vs. OCM indicated by *. P-values are given in Table 1.

### Patch-clamp electrophysiological recording and analysis

Electrophysiological recordings were made from cultures at 17 DIV. Dishes were transferred to a recording chamber mounted on a BX51WI microscope (Olympus, France) and superfused with ACSF containing (in mM): 124 NaCl, 2.5 KCl, 26 NaHCO_3_, 1 NaH_2_PO_4_, 2 CaCl_2_, 2 MgCl_2_, and 11 glucose, bubbled with 5% CO_2_ in O_2_ (pH 7.3, 305-315 mOsm/L). Temperature was kept at 34° C. Recordings were made with glass pipettes pulled using a Brown–Flaming electrode puller (Sutter Instruments) from borosilicate glass of external diameter 1.5 mm (Clark Capillary Glass, Harvard Apparatus). Pipette resistance was 6 MΩ when filled with a solution containing (in mM): 135 K-gluconate, 1.2 KCl, 10 HEPES, 0.2 ethylene glycol tetraacetic acid (EGTA), 2 MgCl_2_, 4 MgATP, 0.4 Tris-GTP, 10 Na_2_-phosphocreatine and 2.7 biocytin. RNase inhibitor was added (5μl in 1ml) when harvesting cell contents. Pipette solution pH was adjusted to 7.3 with KOH and the osmolarity was 290 mOsm. Whole-cell current-clamp recordings were made using a MultiClamp 700B amplifier and pCLAMP software (Molecular Devices). Potential signals were filtered at 6 kHz and digitized at 20–50 kHz.

**Table 1:**
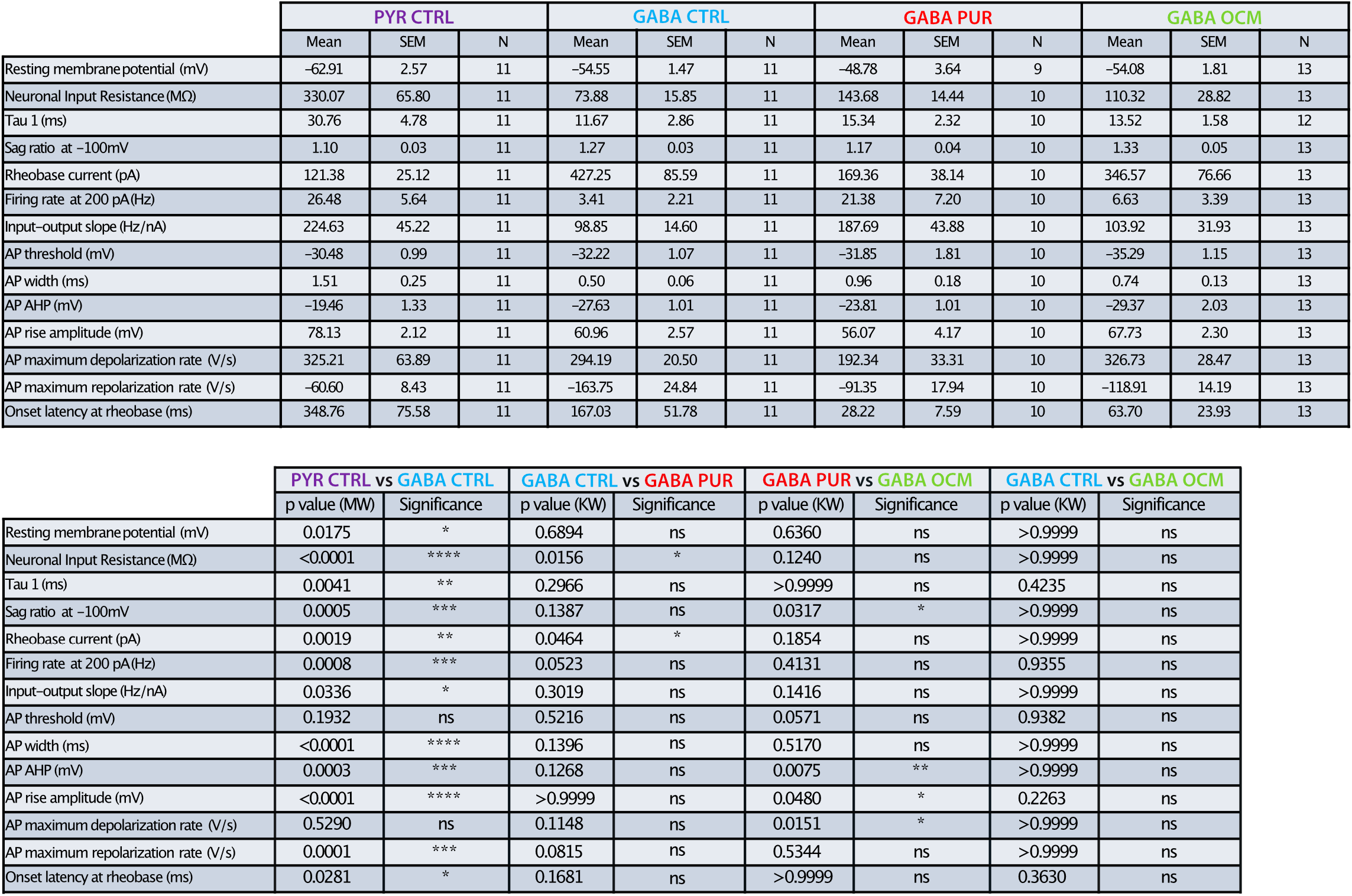
Electrophysiological properties of pyramidal and GABAergic neurons. Top, mean values ± SEM and number of cells for each parameter and culture condition. Bottom, p-values and significance levels from Mann-Whitney (MW) tests (PYR CTRL vs. GABA CTRL) and from Kruskal-Wallis (KW) and Dunn’s multiple comparison *post hoc* test (GABA CTRL vs. PUR vs. OCM).

|  | PYR CTRL |  |  | GABA CTRL |  |  | GABA PUR |  |  | GABA OCM |  |  |
| --- | --- | --- | --- | --- | --- | --- | --- | --- | --- | --- | --- | --- |
|  | Mean | SEM | N | Mean | SEM | N | Mean | SEM | N | Mean | SEM | N |
| Resting membrane potential (mV) | -62.91 | 2.57 | 11 | -54.55 | 1.47 | 11 | -48.78 | 3.64 | 9 | -54.08 | 1.81 | 13 |
| Neuronal Input Resistance (MΩ) | 330.07 | 65.80 | 11 | 73.88 | 15.85 | 11 | 143.68 | 14.44 | 10 | 110.32 | 28.82 | 13 |
| Tau 1 (ms) | 30.76 | 4.78 | 11 | 11.67 | 2.86 | 11 | 15.34 | 2.32 | 10 | 13.52 | 1.58 | 12 |
| Sag ratio at -100mV | 1.10 | 0.03 | 11 | 1.27 | 0.03 | 11 | 1.17 | 0.04 | 10 | 1.33 | 0.05 | 13 |
| Rheobase current (pA) | 121.38 | 25.12 | 11 | 427.25 | 85.59 | 11 | 169.36 | 38.14 | 10 | 346.57 | 76.66 | 13 |
| Firing rate at 200 pA (Hz) | 26.48 | 5.64 | 11 | 3.41 | 2.21 | 11 | 21.38 | 7.20 | 10 | 6.63 | 3.39 | 13 |
| Input-output slope (Hz/nA) | 224.63 | 45.22 | 11 | 98.85 | 14.60 | 11 | 187.69 | 43.88 | 10 | 103.92 | 31.93 | 13 |
| AP threshold (mV) | -30.48 | 0.99 | 11 | -32.22 | 1.07 | 11 | -31.85 | 1.81 | 10 | -35.29 | 1.15 | 13 |
| AP width (ms) | 1.51 | 0.25 | 11 | 0.50 | 0.06 | 11 | 0.96 | 0.18 | 10 | 0.74 | 0.13 | 13 |
| AP AHP (mV) | -19.46 | 1.33 | 11 | -27.63 | 1.01 | 11 | -23.81 | 1.01 | 10 | -29.37 | 2.03 | 13 |
| AP rise amplitude (mV) | 78.13 | 2.12 | 11 | 60.96 | 2.57 | 11 | 56.07 | 4.17 | 10 | 67.73 | 2.30 | 13 |
| AP maximum depolarization rate (V/s) | 325.21 | 63.89 | 11 | 294.19 | 20.50 | 11 | 192.34 | 33.31 | 10 | 326.73 | 28.47 | 13 |
| AP maximum repolarization rate (V/s) | -60.60 | 8.43 | 11 | -163.75 | 24.84 | 11 | -91.35 | 17.94 | 10 | -118.91 | 14.19 | 13 |
| Onset latency at rheobase (ms) | 348.76 | 75.58 | 11 | 167.03 | 51.78 | 11 | 28.22 | 7.59 | 10 | 63.70 | 23.93 | 13 |

Table 1
|  | PYR CTRL vs GABA CTRL |  | GABA CTRL vs GABA PUR |  | GABA PUR vs GABA OCM |  | GABA CTRL vs GABA OCM |  |
| --- | --- | --- | --- | --- | --- | --- | --- | --- |
|  | p value (MW) | Significance | p value (KW) | Significance | p value (KW) | Significance | p value (KW) | Significance |
| Resting membrane potential (mV) | 0.0175 | * | 0.6894 | ns | 0.6360 | ns | >0.9999 | ns |
| Neuronal Input Resistance (MΩ) | <0.0001 | **** | 0.0156 | * | 0.1240 | ns | >0.9999 | ns |
| Tau 1 (ms) | 0.0041 | ** | 0.2966 | ns | >0.9999 | ns | 0.4235 | ns |
| Sag ratio at -100mV | 0.0005 | *** | 0.1387 | ns | 0.0317 | * | >0.9999 | ns |
| Rheobase current (pA) | 0.0019 | ** | 0.0464 | * | 0.1854 | ns | >0.9999 | ns |
| Firing rate at 200 pA (Hz) | 0.0008 | *** | 0.0523 | ns | 0.4131 | ns | 0.9355 | ns |
| Input-output slope (Hz/nA) | 0.0336 | * | 0.3019 | ns | 0.1416 | ns | >0.9999 | ns |
| AP threshold (mV) | 0.1932 | ns | 0.5216 | ns | 0.0571 | ns | 0.9382 | ns |
| AP width (ms) | <0.0001 | **** | 0.1396 | ns | 0.5170 | ns | >0.9999 | ns |
| AP AHP (mV) | 0.0003 | *** | 0.1268 | ns | 0.0075 | ** | >0.9999 | ns |
| AP rise amplitude (mV) | <0.0001 | **** | >0.9999 | ns | 0.0480 | * | 0.2263 | ns |
| AP maximum depolarization rate (V/s) | 0.5290 | ns | 0.1148 | ns | 0.0151 | * | >0.9999 | ns |
| AP maximum repolarization rate (V/s) | 0.0001 | *** | 0.0815 | ns | 0.5344 | ns | >0.9999 | ns |
| Onset latency at rheobase (ms) | 0.0281 | * | 0.1681 | ns | >0.9999 | ns | 0.3630 | ns |

Recordings in the whole-cell current clamp configuration were made from fluorescent GABAergic neurons and non-fluorescent pyramidal shaped neurons in cultures prepared from VGAT-Venus Wistar rats. Excitatory postsynaptic potential (EPSP) and action potential (AP) frequencies, membrane potential and input resistance were measured at resting potential. Responses to families of hyperpolarizing and depolarizing current steps of duration 800 ms were recorded from holding potentials near −60 mV. Current intensities were manually adjusted to induce a maximal hyperpolarization close to −100 mV. Incremental positive steps of +1/10 of that value were then applied until the cell was depolarized above rheobase several times. After recording electrical data for ~10 min, neuronal contents were aspirating into the glass electrode tip. They were extracted into a tube containing 3.5 μl of lysis buffer with RNase inhibitor as a first step to prepare a library of neuronal total RNA (Qiu et al., 2012). Electrophysiological signals were analyzed with AxographX and routines written in MATLAB (The Mathwork; Huang et al. 2017). EPSP and AP frequencies were measured from baseline records of duration at least 3000 ms. Sup. Fig1 shows procedures used to measure active and passive membrane parameters.

### Reconstruction of neuronal morphology

Pipettes used for patch-clamp recordings included 2.7 mM biocytin. Cultures containing filled cells were fixed at 17 DIV with PFA4% (diluted in PBS 1X; pH 7.2) for 10 min at room temperature (RT). Coverslips were washed three times with PBS 1X and blocked with 5% normal goat serum containing 0.1% Triton for 15 min at RT. Biocytin-filled cells and their axon initial segments were visualized. Cultures were incubated with anti-Neurofascin (1:100, ab31457, Abcam) for 2 hours at RT. After three PBS rinses, they were incubated with Streptavidine-Alexa 488 (ThermoFisher Scientific) to visualize biocytin-filled neurons and anti-rabbit-Alexa 594 (1:1000, ThermoFisher Scientific) for Neurofascin for 1 hour at RT. Stained cultures were mounted with Fluoromount-G.

Images of stained cells were acquired on a upright spinning disk microscope (Intelligent Imaging Innovations, Inc) using a 20x glycerol immersion objective (NA 1.0), a CSU-W1 spinning disk head (Yokogawa) and a sCMOS ORCA-Flash4.0 camera (Hamamatsu). Multiple tile regions each with Z step series of separation 1.1 μm were acquired for each cell. Tile scans were stitched using Fiji software with BigStitcher plugin. Neuronal arborizations were drawn with the semi-automatic filament tracer tool of IMARIS software (Bitplane). The axon was identified from Neurofascin immunostaining of its initial segment. Axonal and dendritic lengths and data for Sholl analyses were derived by the IMARIS software.

### Statistical analysis of electrophysiological properties and morphology

Statistical analyses were performed using GraphPad Prism version 7.0. Experiments for each condition were carried out in at least 3 independent cultures from at least 3 different litters. Electrophysiological parameters in Figs. 1 and 2 were compared using the Mann-Whitney test (for PYR CTRL vs. GABA CTRL) and using Kruskal Wallis and Dunn’s multiple comparison *post hoc* test (for GABA CTRL vs. GABA PUR, GABA PUR vs. GABA OCM and GABA CTRL vs. GABA OCM). P-values are given in Table 1.

**Figure 2:**
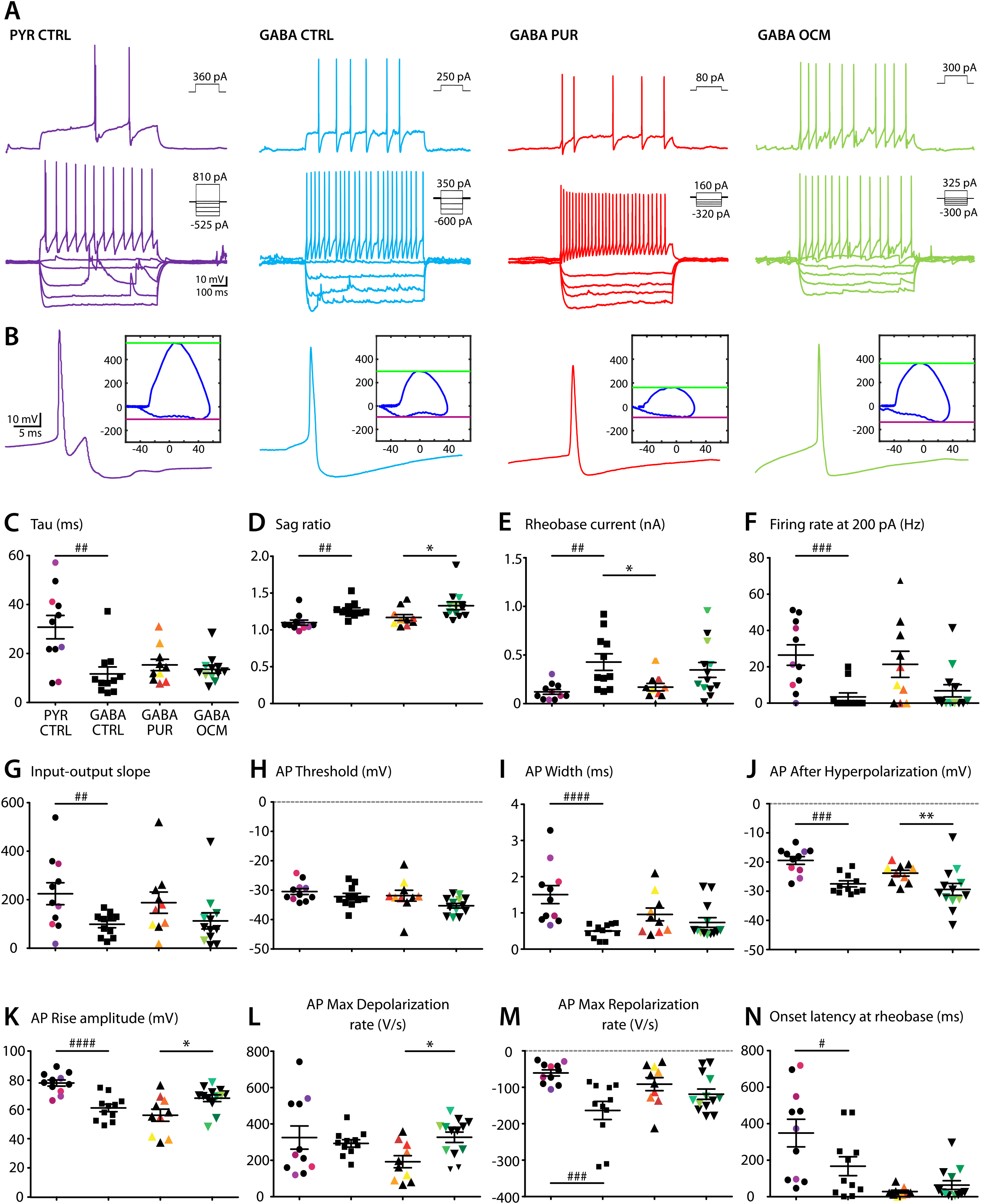
Glial factors affect the electrophysiological properties of GABAergic neurons. (A) Voltage responses to depolarizing and hyperpolarizing current steps recorded from representative neurons at 17 DIV in the different conditions. Top, action potentials initiated at rheobase; bottom, current intensities as shown in insets. (B) Action potential waveforms and phase plots (Y axis, dV/dt (V/s); X axis, membrane potential (mV). Green line, maximum depolarization rate; blue line, maximum repolarization rate. (C) to (N), Effects of culture conditions (PYR CTRL, GABA CRTL, GABA PUR and GABA OCM) on 12 parameters characterizing neuronal intrinsic properties. Each symbol is one neuron. Color symbols correspond to sequenced cells as in Fig. 1 F, G. Significance levels indicated as in Fig. 1, *i.e.* # for Mann-Whitney test and * for Kruskal-Wallis and Dunn’s *post hoc* test. Parameters measured as shown in Sup. Fig. 1. P-values given in Table 1.

Axonal and dendritic lengths were compared (Fig. 3) using Kruskal Wallis and Dunn’s multiple comparison *post hoc* test (for GABA CTRL vs. GABA PUR and GABA PUR vs. GABA OCM and GABA CTRL vs. GABA OCM). Dendritic arborizations were assessed with Sholl analysis which measures the number of dendrites which intersect circles of increasing distance from the neuronal soma (20 μm increments were used). They were analyzed using a linear mixed-effects model (LMM) with culture condition and radial distance as fixed effects, and the cell identifier number as a random effect to account for the successive measurements over the concentric rings (Wilson et al. 2017). Significance for the main effects of condition, distance and their interaction was then evaluated using ANOVA Type II Wald chi-square tests. Analyses were made with R (R Development Core Team, ver 3.5.1, 2019) and plots were generated with the ggplot2 package (Wickham et al., 2016). LMM was fitted with the function lmer in the lme4 package (Bates et al., 2015). When a factor had a significant effect or when a significant interaction was found between condition and distance, *post-hoc* pairwise comparisons were completed with Tukey’s method. Data were square root transformed before modeling to improve model assumptions of linearity, normality and constant variance of residuals,

**Figure 3:**
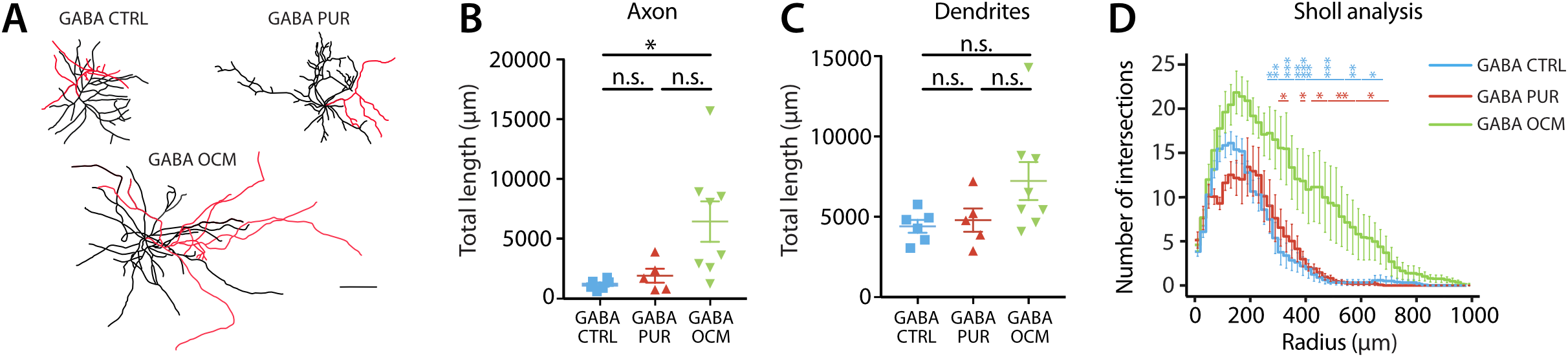
Axo-dendritic morphologies of biocytin filled GABAergic neurons. (A) Reconstruction of biocytin-filled GABAergic neurons at 17 DIV in different culture conditions. Axons shown in red, dendrites in black. Scale bar: 100 μm. (B) Total axonal and (C) dendritic lengths of GABAergic neurons in different conditions (Kruskal-Wallis followed by Dunn’s *post hoc)*. (D) Sholl analysis showing for each condition the number of dendrites intersecting increasing radii (20-1000 μm from the soma at interval 20 μm). The two segmented lines above the plot indicate significant differences between GABA OCM and CTRL (blue) and between GABA OCM and PUR (red). There were no significant differences between GABA CTRL and PUR. LMM and Type II Wald chi-square tests followed by *post-hoc* analyses using Tukey’s method. The mean ± SEM of three independent experiments is shown. (\**p* < 0.05, \*\**p* < *0.01*, *\*\*\*p* < 0.001, n.s. indicates no significance).

Data are presented as mean ± SEM (*i.e.* standard error of the mean). The level of statistical significance was set at p < 0.05 for all tests. Significance is represented as numbers of asterisks: *p < 0.05, **p < 0.01, ***p < 0.001, ****p < 0.0001 and n.s. indicates no significance.

### cDNA synthesis, library preparation and sequencing

mRNA capture, reverse transcription and amplification was achieved using the SMART-Seq v4 ultra low input RNA Kit (Takara, 634891). This kit improves synthesis of the full-length cDNA via a template switching mechanism for synthesis of the second strand cDNA. 5μl of sample was used for hybridization with the 3’smart-seq primer, then poly{T}-primed mRNA was converted to cDNA by reverse transcriptase and PCR amplification was done, according to the kit instructions. Full-length cDNA was then processed with a Nextera XT DNA Library Preparation Kit (Illumina, FC-131-1096). This kit aims to fragment and add adapter sequences onto template DNA with a single tube Nextera XT tagmentation reaction and so generate multiplex sequencing libraries. The resulting indexed paired-end libraries were sequenced by next-generation sequencing (NGS), using NextSeq500 2×75pb (Illumina NextSeq 500 platform) (Sup. Fig2A).

### RNA-seq data analysis

RNA-Seq data analysis was performed by GenoSplice technology (www.genosplice.com). Sequencing, data quality, read distribution (to check for ribosomal contamination for instance), and insert size estimation were done with FastQC, Picard-Tools, Samtools and rseqc. Reads were mapped using STARv2.4.0 (Dobin et al. 2013) on the rn6 Rat genome assembly. The regulation of gene expression was studied as in (Noli et al. 2015). For each gene of the Rat FAST DB v2016_1 annotations, reads aligning on constitutive regions (not prone to alternative splicing) were counted. Normalization and differential gene expression was assessed from these reads using DESeq2 running on R (v.3.2.5, Love et al. 2014). FastQC was used for quality control, sequencing quality per base and sequence, per base sequence and GC content, N content, overrepresented sequences, sequence lengths. Sequence coverage of introns and exons was used to ensure that the sequences derived from mRNA rather than genomic DNA.

From 64 samples, 21 passed these control quality steps. Others were discarded due to sample quality (n=32) or contamination detected from marker expression (n=11) based on the following *GFAP*, *Aquaporin 4, Slc1a2, PDGFRa, MOG, Itgam* (Sup. Fig2 B, D, E).

Genes were considered as expressed if their FPKM (*i.e.* Fragments Per Kilobase Million) value was greater than 98% of the background FPKM value from intergenic regions (Sup. Fig3).

Clustering and heatmap analyses were performed using dist” and “hclust” functions in R, with Euclidean distance and Ward agglomeration method.

Genes were considered to be differentially expressed when the uncorrected p-value ≤ 0.05 and fold-change ≥ 1.5.

Enriched GO terms were assessed, from the identity of differentially expressed genes, with the DAVID Functional Annotation Tool (v6.8), (Huang et al. 2007). GO terms were considered enriched when fold enrichment ≥ 2.0 and uncorrected p-value ≤ 0.05 (1.3 value in GO term graph matches with a p value of 0.05), and at least 2 regulated genes in the pathway/term.

A Pearson correlation-based approach was used to compare single-cell RNA values with electrophysiological parameters for neurons (Fig. 6). The procedure was restricted to genes expressed in at least five samples, with coefficients greater than 0.6 and significant p-values (p<0.05).

## RESULTS

### Oligodendrocyte secreted factors control the electrophysiological properties of hippocampal GABAergic neurons

We first asked how the presence of glial cells or oligodendrocyte secreted factors affected GABAergic neuron phenotype. Hippocampal neuron cultures were prepared from VGAT-Venus rat embryos so that GABAergic interneurons could be identified. Spontaneous activity and active and passive membrane properties were recorded in the current clamp mode from fluorescent GABAergic neurons (n=36). These properties were compared with those of unlabeled pyramidal cells (n=11). We further compared the physiology of GABAergic neurons in control cultures (CTRL) with those in purified cultures lacking glial cells (PUR) and with those in purified cultures supplemented with oligodendrocyte conditioned medium (OCM) (Fig. 1A, B).

Spontaneous synaptic events and spiking activity of recorded cells was quantified at resting potential with no injected current (Fig. 1C-E). In pyramidal cells (PYR), frequencies of excitatory postsynaptic potentials (EPSP) and action potentials (AP) were low in CTRL cultures (PYR CTRL, EPSPs, 7±2 Hz; APs, 0.03±0.01 Hz; n=11). Mean resting potential was −63±3 mV and mean input resistance was 330±66 MΩ (Fig. 1F, G; Table 1). EPSP frequency and AP discharge rate were both higher in GABAergic neurons from the same cultures. Mean EPSP frequency was 70±13 Hz and AP discharge frequency was 0.6±0.5 Hz (GABA CTRL, n=11).

In PUR neuronal cultures, both EPSP frequency and AP discharge by GABAergic neurons were reduced (GABA PUR, EPSPs, 6±2 Hz, n=10; APs, 0.03±0.03 Hz, n=11). This despite a more depolarized mean resting membrane potential (GABA PUR, −49±4 mV vs GABA CTRL, −55±2 mV) and a significantly higher input resistance than in control cultures (GABA PUR, 144±14 MΩ vs GABA CTRL, 74±16 MΩ). Supplementing PUR cultures with oligodendrocyte conditioned medium tended to increase EPSP and AP frequencies towards CTRL levels (GABA OCM, EPSPs, 19±4 Hz, n=13; APs, 0.4±0.03 Hz, n=12). Mean membrane potential and input resistance also reverted towards values in CTRL mixed cultures (GABA OCM, −54±2 mV and 110±29 MΩ). These data suggest that oligodendrocyte secreted factors influence active and passive aspects of the physiological phenotype of GABAergic neurons.

We next asked whether the properties of fluorescent and non-fluorescent cells were consistent with those of GABAergic neurons and pyramidal cells respectively (Pelkey et al. 2017). In control cultures, the mean resting potential of fluorescent neurons was significantly more depolarized (GABA CTRL, −55±1 mV vs. PYR CTRL, −63±3 mV; Fig. 1F), and their input resistance was significantly lower than that of non-fluorescent cells (74±16 MΩ vs. 330±66 MΩ; Fig. 1G). Comparing neuronal responses to depolarizing and hyperpolarizing step current injections (Sup. Fig. 1, Fig. 2) revealed significant differences in the membrane time constant tau (Fig. 2C, GABA CTRL, 12±3 ms vs. PYR CTRL, 31±5 ms), sag ratio, which is linked to the presence of an h-current, (Fig. 2D, 1.27±0.03 vs 1.10±0.03), rheobase current (Fig. 2E, 427±86 pA vs 121±25 pA) and firing rate induced by a 200 pA step current injection (Fig. 2F, 3±2 Hz vs 26±6 Hz). Input-output plots of the number of APs against the injected current had a mean initial slope of 99±15 Hz/pA in fluorescent cells, significantly lower than 224±45 Hz/pA for non-fluorescent cells (Fig. 2G). Action potential width in fluorescent neurons was short, 0.50±0.06 ms (Fig. 2I), as is characteristic of some interneurons, compared to an AP width of 1.51±0.25 ms in non-fluorescent neurons. AP thresholds were similar: −32.2±1.1 mV in fluorescent cells (Fig. 2H) and −30.5±0.1 mV in non-fluorescent cells. AP rising amplitude (Fig. 2K) was 61±3 mV in fluorescent cells, significantly lower than a value of 78±2 mV in non-fluorescent cells. Maximum depolarization and repolarization rates were 294±21 V/s and - 164±25 V/s respectively in fluorescent cells, compared to 325±64 V/s (Fig. 2L) and −61±8 V/s (Fig. 2M) in non-fluorescent neurons. Action potential after hyper-polarizations (AHP) were larger in fluorescent cells at −28±1 mV compared to −19±1 mV (Fig. 2J). The latency to the first AP at rheobase was significantly shorter, 167±52 ms compared to 349±76 ms (Fig. 2N). Overall these data confirm that fluorescent neurons from VGAT-Venus animals correspond to GABAergic neurons, and non-fluorescent cells to pyramidal cells. Table 1 summarizes these physiological data and provides statistical support for comparisons.

Our next objective was to compare electrophysiological phenotypes for GABAergic neurons in control conditions (CTRL), in purified cultures (PUR) with no glial cells and in PUR cultures supplemented with oligodendrocyte conditioned medium (OCM). We found nodal proteins were clustered on GABAergic axons in CTRL but not in PUR cultures (not shown). Adding OCM to PUR cultures induced prenode formation as previously shown (Freeman et al. 2015; Dubessy et al. 2019). We also found some electrophysiological parameters of GABAergic neurons changed when glial cells were absent, and were partially restored by OCM treatment (Fig. 1, 2 and Table 1). The mean resting membrane potential of GABAergic neurons did not change significantly (Fig. 1F, CTRL, −55±2 mV; PUR, −49±4 mV; OCM, −54±2 mV). Input resistance increased in PUR cultures (Fig. 1G, PUR, 144±14 MΩ, GABA CTRL, 74±16 MΩ) and was reduced back towards control values by OCM addition (110±29 MΩ). Tau did not change significantly (Fig. 2C, PUR, 15±2 ms; OCM, 14±2 ms). Sag ratio decreased significantly in PUR (Fig. 2D, PUR, 1.17±0.04; OCM, 1.33±0.05), as did the rheobase current (2E, PUR, 169±38 pA; OCM, 347±77 pA. Mean firing rate induced by 200 pA step current injection increased in PUR (Fig. 2F, PUR, 21±7 Hz; OCM, 7±3 Hz). The slope of input-output curves did not change (Fig. 2G, PUR, 188±44 Hz/pA; OCM, 104±32 Hz/pA). Mean AP threshold (Fig. 2H) was −31.9±1.8 mV in PUR, and −35.3±1.2 mV in OCM. AP width increased in PUR cultures (Fig. 2I, PUR, 0.96+0.18 ms; OCM, 0.74±0.13 ms), while AHP amplitude decreased (Fig. 2J, PUR, −24±1 mV; OCM, −29±2 mV). AP rising amplitude was significantly higher in OCM than in PUR cultures (Fig. 2K, PUR, 56±4 mV; OCM, 68±2 mV). Both the AP depolarization rate (Fig. 2L, PUR, 192±33 V/s; OCM, 327±28 V/s) and maximal repolarization rate were slower (Fig. 2M, PUR, −91±18 V/s; OCM, −119±14 V/s) in PUR culture compared to CTRL and OCM cultures. AP onset latency at rheobase was shorter in PUR conditions than in CTRL (Fig. 2N, PUR, 28±8 ms; OCM, 64±24 ms).

These data show that factors of the OCM have significant effects on the input resistance, rheobase current, and AP and AHP amplitudes of GABAergic neurons. These effects partially reverse changes induced by switching from a mixed culture of neurons, oligodendrocytes, and other glial cells, to a purified neuronal culture.

### Impact of OCM on hippocampal GABAergic neuron morphology

We next asked whether factors secreted by oligodendrocyte exert trophic effects on GABAergic neuron anatomy. Biocytin-filled GABAergic neurons from CTRL, PUR, and OCM cultures were reconstructed and axonal and dendritic lengths were measured (Fig. 3A-C). The absence of glial cells had no significant effect on total axonal length (CTRL, 1179 ± 174 μm, n=6; PUR, 1929 ± 583 μm, n=5; Fig. 3B) or total dendritic length (CTRL, 4405 ± 401 μm, n=6; PUR, 4794 ± 725 μm, n=5; Fig. 3C). Scholl analysis of dendritic arbors revealed slight differences (Fig. 3D). However, in OCM cultures, total axonal length (OCM, 6451 ± 1695 μm, n=8) was significantly increased compared to CTRL (p=0.0148) and mid-dendritic arbors were significantly more complex at distances of 260-700 μm from the soma compared to CTRL and PUR (Fig. 3B, D). Thus oligodendrocyte derived factors exert effects on axons and dendrites of GABAergic neurons, which were not apparent on switching from CTRL to PUR cultures.

### Single-cell transcriptomic analysis of electrophysiologically characterized neurons

We used single cell RNA-sequencing (scRNA-seq) of the cytoplasmic contents of GABAergic neurons to pursue the identity of protein targets underlying changes in their phenotype induced by OCM. When patch-clamp recordings were completed, the same pipette was used to harvest neuronal cytosolic contents (n=64). Sup. Fig. 2A, B illustrates the experimental workflow and bioinformatics pipeline. We noted a strong correlation (R=0.79) between the number of expressed genes and the number of aligned reads suggesting that increasing reads could improve transcript detection (Fig. 4-1C). RNA-seq data was filtered to exclude samples of poor RNA quality where fewer than 2000 transcripts were detected. We also excluded samples which expressed glial cell specific genes, such as *Mog* or *Gfap*, as likely contaminated. Samples from 21 neurons passed these two controls.

**Figure 4:**
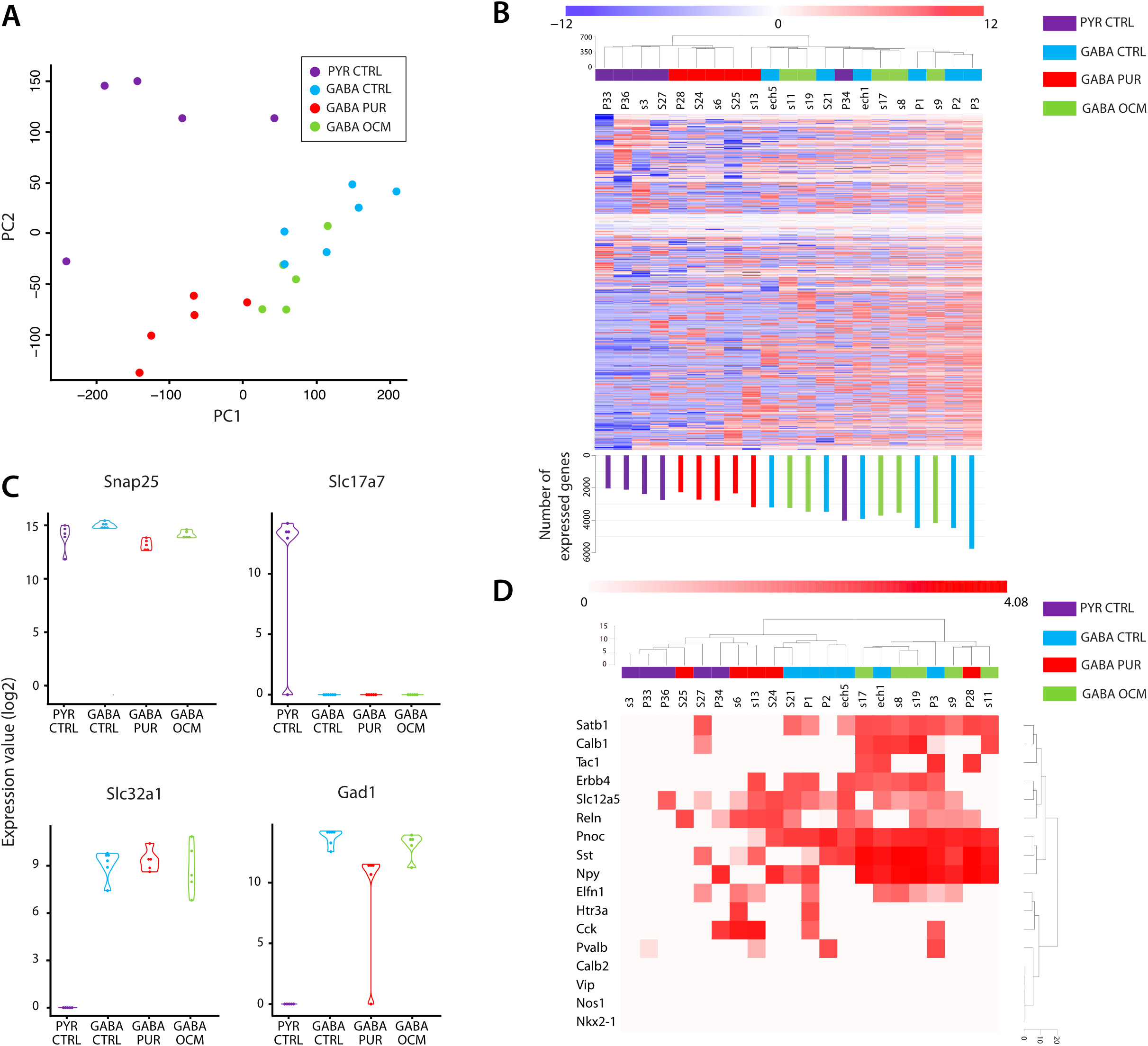
Gene expression analysis of the content of pyramidal or GABAergic neuron cyto-plasm in the presence or absence of glial factors. (A) Principal component analysis showing clustering of validated neurons by culture conditions on single-cell data. PC1 and PC2 explain 14.9 % and 6.6 % of the variance, respectively. (B) Heatmap summarizing unsupervised analysis of mRNAs from validated neurons in different culture conditions. Rows represent genes and columns different neurons. Colors indicate gene expression (blue, low – red, high) normalized by row. Barplots (below) show the number of genes expressed by each neuron. 5391 genes (out of 35 152) are represented here and only genes detected in at least 20% of neurons are included. C) Expression levels for markers for neurons (*Snap25*), excitatory neurons (*Slc17a7*) and GABAergic neurons (*Slc32a1* and *Gad1*) in different culture conditions. (D) Differential expression of genes associated with subtypes of GABAergic neuron. Rows represent gene expression and columns represent different neurons under different culture conditions. Color intensity represents gene expression level. In A-D, data from pyramidal neurons under CTRL conditions are shown in purple, GABAergic neurons in CTRL conditions in blue, GABAergic neurons in purified neuron cultures in red and GABAergic neurons in purified cultures treated with OCM in green.

Principal component analysis (PCA) of transcriptomic profiles captured 22% of explained variance with the first two principal components, PC1 and PC2 (Fig. 4A). Pyramidal cells and GABAergic neurons were clearly segregated in this two-dimensional space. GABAergic neurons from PUR cultures were separated from GABAergic neurons in CTRL cultures. Transcriptomes of GABAergic neurons from OCM cultures overlapped with those of CTRL cultures.

We investigated transcriptomic differences between hippocampal neuron types and the transcriptional response to glial factors in GABAergic neurons. The heat map of Fig. 4B shows in red up-regulated genes and in blue genes that were down-regulated with respect to mean expression in all three conditions. High, deep sequencing detected up to 5,700 expressed genes in single neurons. Hierarchical clustering of samples from different cells in the heat map segregated 4 out of 5 pyramidal neurons from GABAergic neurons. GABAergic neurons from purified cultures (PUR) were also grouped together in the dendrogram while GABAergic neurons sampled in CTRL and OCM cultures overlapped in a large branch (Fig. 4B).

As a step towards validation of these results, we searched for the presence of known pyramidal and GABAergic neuron markers in genes from different samples. All samples expressed the neuronal marker *Snap25*. Only samples from non-fluorescent pyramidal cells expressed the vesicular glutamate transporter1 (vGlut1, *Slc17a7)*, while the vesicular GABA transporter (VGAT; *Slc32a1*) and GAD67 (*Gad1*) were only detected in samples obtained from fluorescent GABAergic neurons (Fig. 4C). Subclasses of hippocampal GABAergic neurons express neuropeptides and Ca^2+^-binding proteins. Searching for neuropeptide (Somatostatin (*Sst*), Neuropeptide Y (*Npy*), Cholecystokinin (*Cck*), Vasoactive intestinal peptide (*Vip*), Protachykinin (*Tac1*), Prepronociceptin (*Pnoc*)) and Ca^2+^-binding protein (Parvalbumin (*Pvalb*), Calbindin-1 (*Calb1*), Calbindin-2 (*Calb2*) genes revealed a diversity of expression. Most GABAergic neurons expressed *Pnoc (87%)*, *Sst (81%)* and *Npy* (57%). Samples from 56% of cells expressed both *Sst* and *Npy*, or combinations of *Sst* and *Calb1* (37%) and/or *Sst* and *Cck* (25%) and/or *Sst* and *Pvalb* (19%, Fig. 4D). GABAergic neurons expressing *Pnoc, Npy* and *Sst* neuropeptide genes were found in all culture conditions (CTRL, PUR and OCM). *Calb-1* was less frequently expressed in PUR cultures. Genes for other interneuron markers detected in GABAergic neurons included the transcription factors *Satb1* and *Nkx-2.1*, the post-synaptic protein *Elfn1*, the serotonin receptor *Htr3a*, KCC2 the potassium chloride cotransporter 2, *Slc12a5*, the kinase *Erbb4* and the protease reelin, *Reln*. We note that some molecular markers examined here are not entirely specific to GABAergic neurons, and were also detected in samples from 1 or 2 pyramidal cells (*Slc12a5*, *Calb1*, *Satb1, Reln, Sst, Npy, Elfn1 and Cck.*). Furthermore, no samples from either pyramidal or GABAergic neurons expressed *Vip, Calb2* or *Nos1* (Fig. 4D).

### Ion channel and transmitter receptor gene expression in single hippocampal neurons

The cellular and synaptic physiology of neurons depends on the expression of genes coding for ion channels, transporters and neurotransmitter receptors. We examined quantitative expression of these genes in samples from each neuron in our RNAseq data set (Fig. 5A, B). Different neurons expressed different levels of *Atp1a, b* genes coding for Na^+^/K^+^-transporting ATPase subunits and *Clcn3-7* genes coding for Cl^-^/H^+^ exchangers, which contribute to stabilize membrane potential. The depolarization phase of action potentials is due to opening of voltage-gated sodium channels, which consist of an α-subunit forming pore (*Scn1a-9a*) and auxiliary ß subunits *(Scn1b-4b).* Almost all neurons expressed high levels of *Scn2a* (encodes Nav1.2). In contrast, only GABAergic neurons in CTRL cultures with glial cells or with oligodendroglial factors (OCM) expressed *Scn1a* (Nav1.1) and few expressed *Scn8a* (Nav1.6). The Na^+^ channel modifier 1 (*Scnm1)*, which governs alternative splicing of pre-mRNAs, was detected in samples from some GABAergic neurons but not from pyramidal cells.

**Figure 5:**
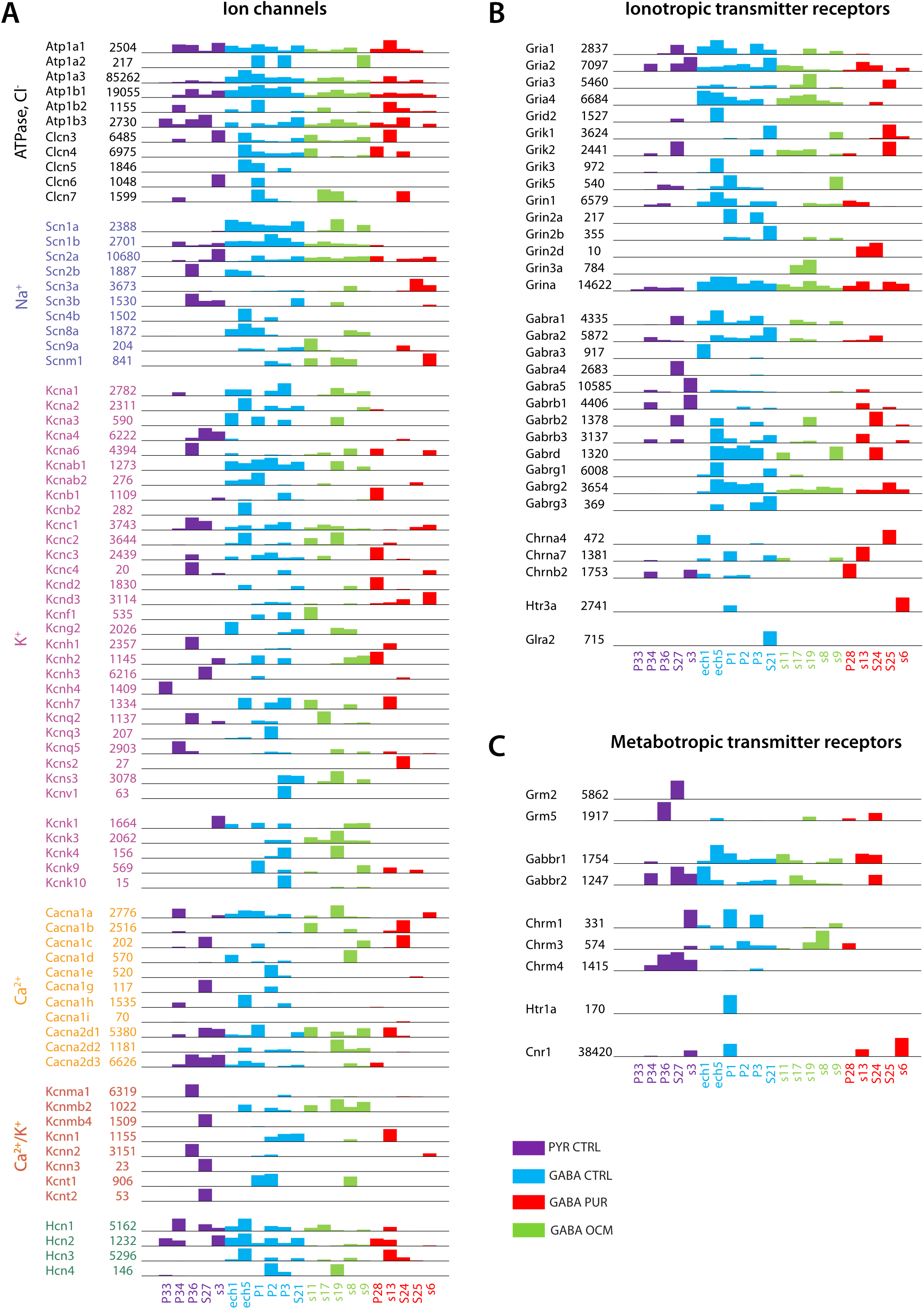
Cell-type specific expression of mRNAs coding for ion channel and receptors. (A) Ion channel, (B) ionotropic transmitter receptor and (C) metabotropic transmitter receptor mRNAs in different culture conditions. Maximal expression from all samples is indicated. Number of reads normalized from pyramidal neurons in CTRL conditions is shown in purple, GABAergic neurons in CTRL conditions in blue, GABAergic neurons in purified cultures in red and GABAergic neurons in purified cultures treated with OCM in green.

Various potassium channels are crucial for action potential repolarization, generate the AHP and contribute to maintenance of membrane potential. Voltage-gated K^+^ channels differ in structure, biophysics and pharmacology from voltage independent, two-pore-domain (K2P) channels which support leak-type K^+^ conductances. We found that distinct neurons express specific combinations of K^+^ channel *∝* (*Kcna-v*) and auxiliary subunits (*Kcnab1-2*) as well as K2P channels (*Kcnk1-10*). We also examined Ca^2+^ channels, Ca^2+^-activated K^+^ channels and hyperpolarization-activated, cyclic nucleotide gated, K^+^/Na^+^ permeable ‘h’ channels (HCN).

Fig. 5B shows genes encoding ionotropic glutamate receptors, expressed at excitatory synapses, and including AMPA receptors (*Gria1-4*), NMDA receptors (*Grin1-3*) and kaand k receptors (*Grik1-5*). Genes encoding GABA_A_ receptors (*Gabra-g*), which mediate fast inhibitory neurotransmission and are assembled as heteropentameric chloride channels, are also indicated as are detected genes which code for subunits of nicotinic cholinergic receptors (*Chrna-b)*, serotonin receptor *(Htr3a)* and glycine receptor *(Glra2).* Fig. 5C shows genes coding for G-protein linked receptors including metabotropic glutamate *(Grm2, 5)* and GABA_B_ *(Gabbr1, 2)* receptors.

### Correlation-based approach

These scPatch-seq data permit quantitative assays of transcriptomic features. We attempted to relate them to intrinsic neuronal electrophysiology by searching for correlations between values from transcriptomic samples and different electrophysiological parameters. Fig. 6 plots genes coding for ion channels, transporters and synaptic receptors for which a correlation (p-value <0.05) with electrophysiological parameters was detected. The analysis was based on all neurons with complete electrical and transcriptomic datasets and on cells from all culture conditions. Genes were clustered based on their relations with electrophysiological parameters. One cluster coding for α1, ß1 and α3 subunits of the Na^+^, K^+^ -ATPase (*Atp1a1*, *Atp1b1* and *Atp1a3*), was correlated with neuronal input resistance, *Atp1b1* was also correlated with time constant, tau and AP width. The Na^+^ channel subunit *Scn2a* (coding for Nav1.2) was correlated with the time constant tau and with AP threshold, neuronal input resistance and AP width. A larger cluster, consisting of several K-channel linked genes (*Kcnc2*, coding for Kv3.2; *Kcnk3*, Task-1; *Kcnip1*, K^+^ channel modulatory protein), as well as two Zn transporters (*Slc30a3*, ZnT3; *Slc30a4*, ZnT4) was strongly correlated with the Sag ratio, the AP threshold and the after-hyperpolarization amplitude (AP AHP).

**Figure 6:**
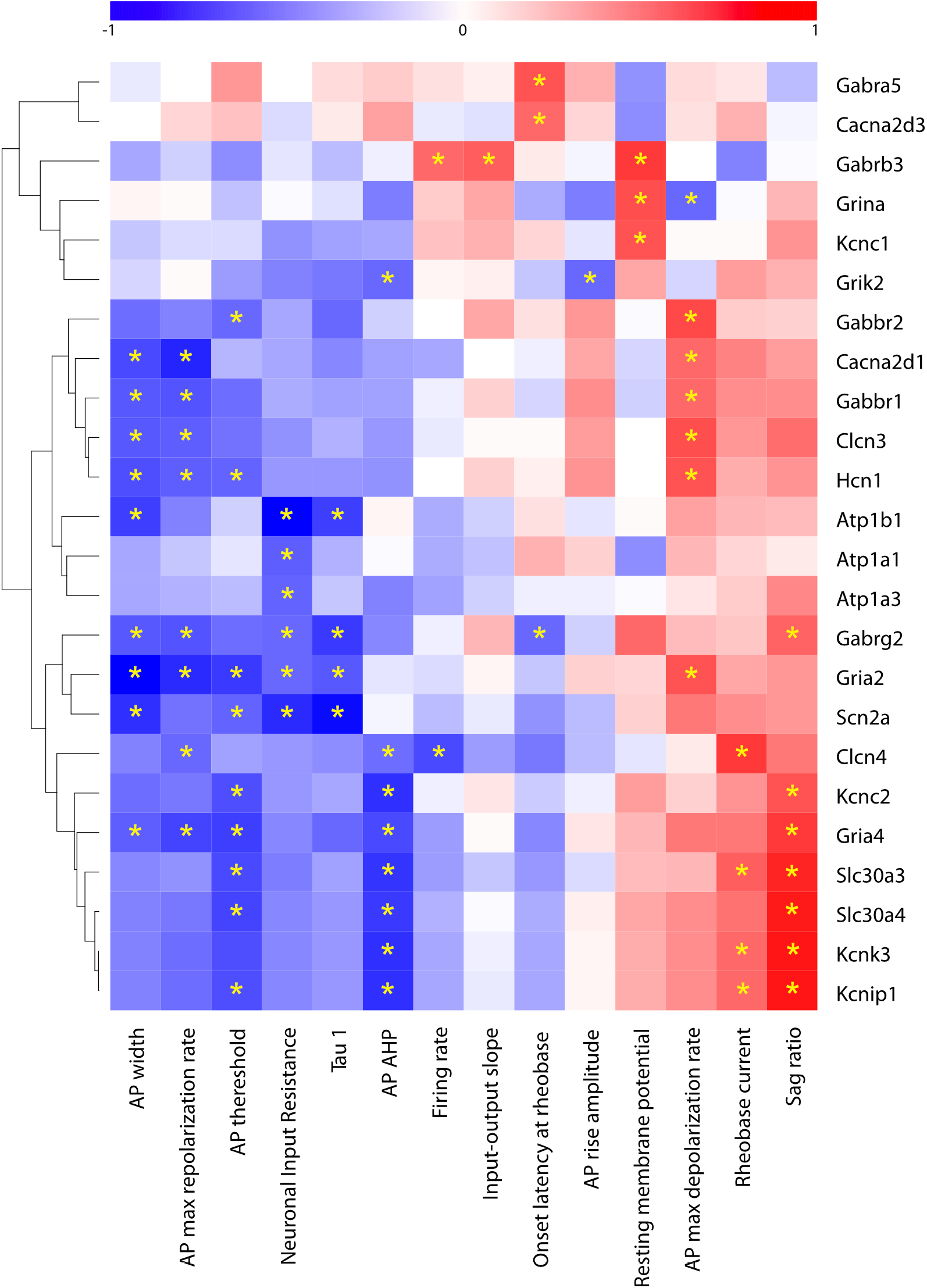
Pearson correlation between scRNASeq data and electrophysiological parameters. Ion channel and synapse related genes significantly correlated (p<0.05) with at least one parameter are shown. Color intensity represents correlation coefficient. Blue indicates negative correlation; expression decreases as the parameter increases. Red indicates positive correlation; both increase together. Significant positive or negative correlations with coefficient > 0.6 and p-value <0.05 are marked with a yellow star.

### Transcriptomic patterns across groups of neurons in relation to biological processes

Gene Ontology (GO) analysis let us estimate biological processes underlying differential expression of groups of genes in different neurons or in different culture conditions. For instance, 326 genes were differentially expressed in GABAergic neurons and pyramidal neurons in CTRL cultures (Fig. 7A). GO process terms derived from the identity of the genes were related to synaptic transmission, organization and transmitter transport as well as cortical development. Comparing GABAergic neurons in CTRL and PUR cultures we found 219 genes were differentially expressed, mostly down-regulated in PUR conditions (Fig. 7B). Inversely, comparing GABAergic neurons in PUR and OCM cultures, 192 genes were differentially expressed, mostly up-regulated in OCM conditions (Fig. 7C). Many genes were down-regulated in one comparison and up-regulated in the other. GO process terms identified from their identity regulate neuronal action potential, synapse assembly, Ca^2+^-dependent exocytosis, protein phosphorylation, kinase signaling and cell division (Fig. 7B, C). GO-analysis further suggested that genes coding for proteins involved in transmembrane transport of K^+^ were up-regulated by OCM treatment (Fig. 7C).

**Figure 7:**
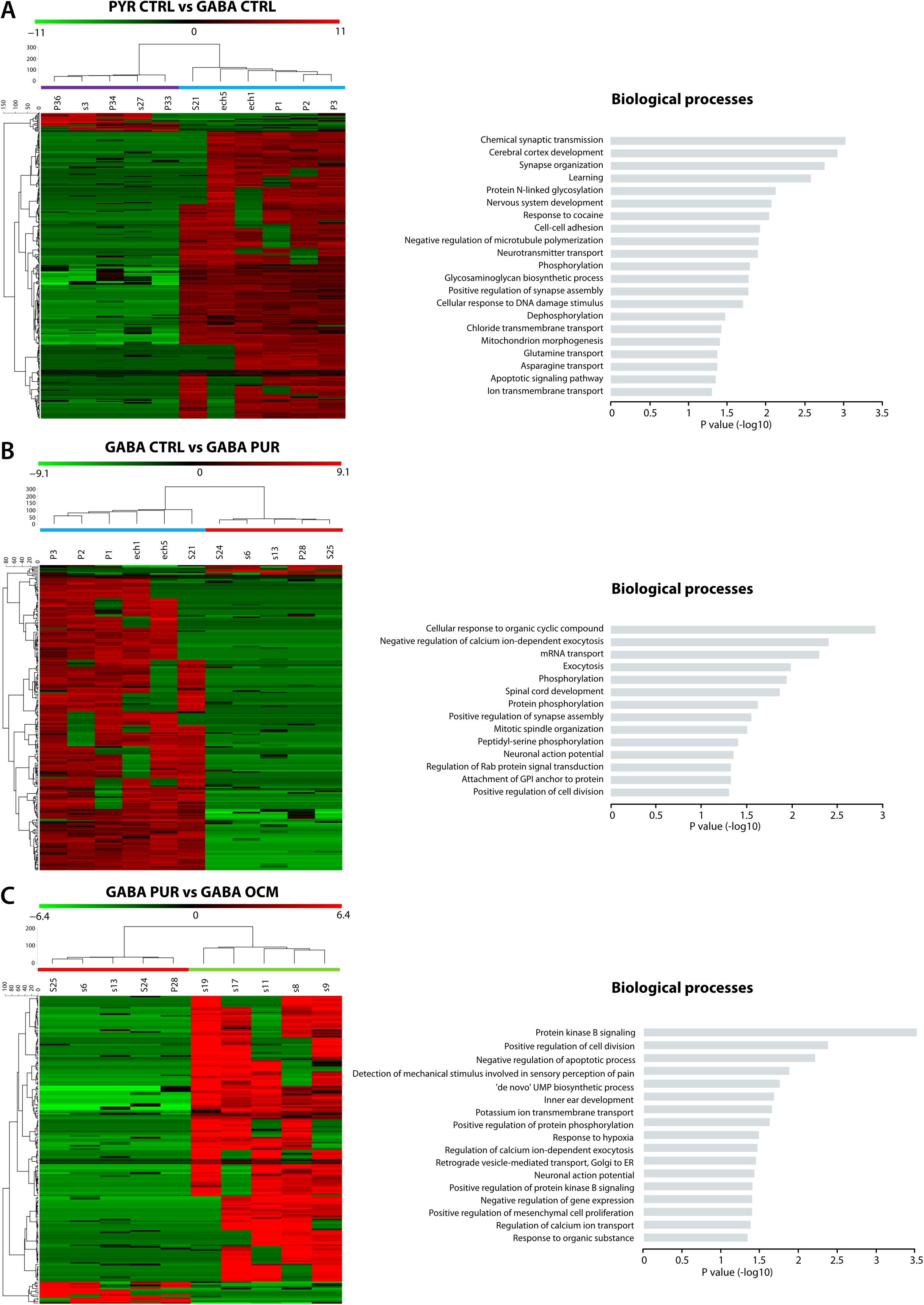
Heatmap of genes expressed differentially between conditions. (A) Differences between pyramidal and GABAergic neurons in CTRL culture conditions. (B) Differences between GABAergic neurons in CTRL and PUR cultures. (C) Effects of OCM on GABAergic neurons in PUR cultures. Rows represent genes and columns represent neurons in different conditions. Color intensity is mean centered expression. At the right, gene ontology analysis of biological processes for differentially expressed genes. Differences between pyramidal and GABAergic neurons in CTRL conditions (upper). Differences between GABAergic neurons in PUR cultures and CTRL. Effects of OCM on GABAergic neurons in PUR cultures (lower).

Further insights into processes affected by oligodendrocyte factors were obtained by grouping genes with similar profiles of changes. RNA-seq data from GABAergic neurons in different culture conditions was normalized with respect to expression of each gene in pyramidal cells (Fig. 8A, B and Sup. Fig4). A total of 241 genes were identified as differentially expressed between pyramidal and GABAergic neurons in CTRL cultures and were mostly insensitive to PUR or OCM culture conditions (Sup. Fig4).

**Figure 8:**
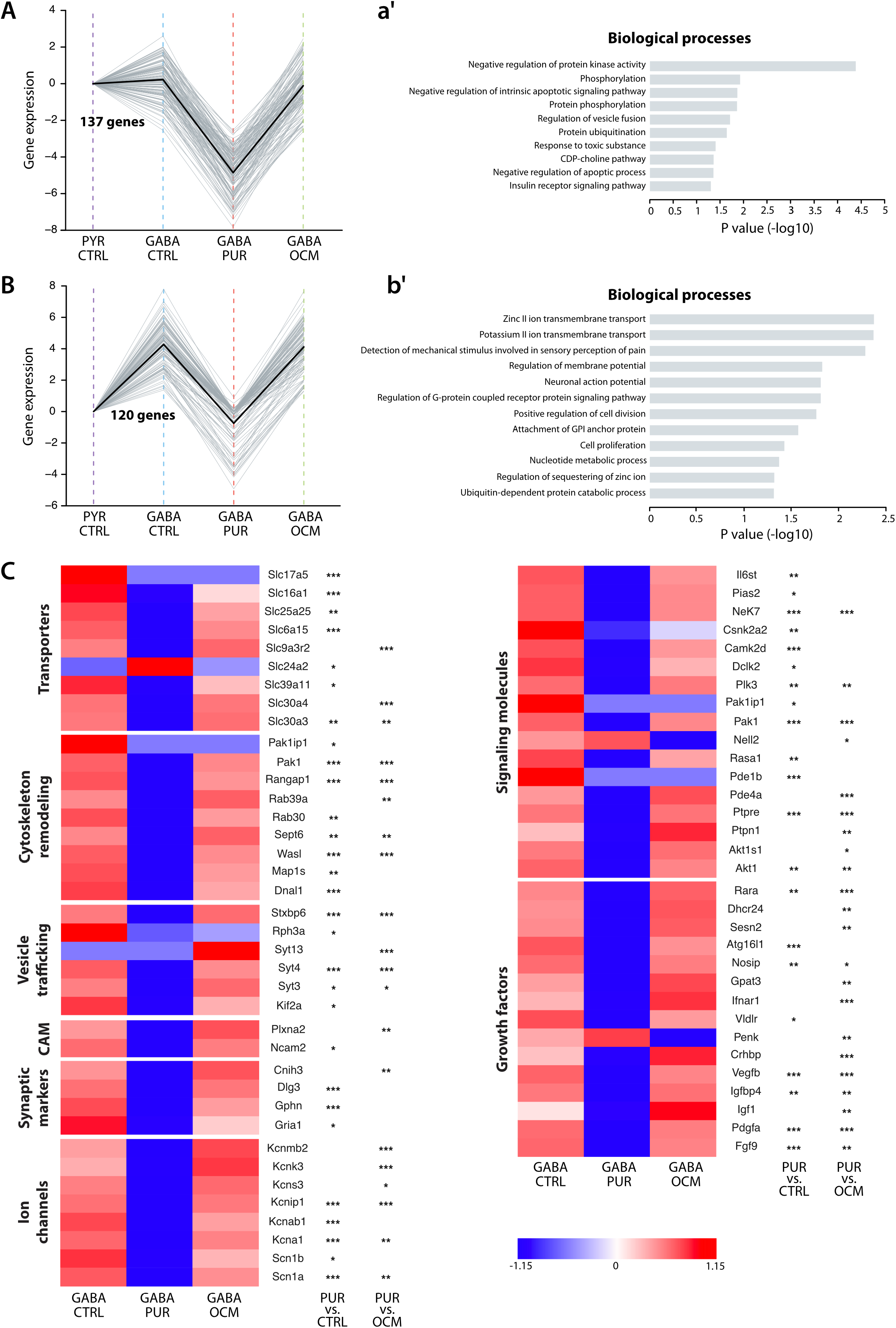
Gene ontology analysis of regulated genes. (A, B) Expression within clusters with a similar expression pattern in different culture conditions. Gene expression was normalized to that of pyramidal cells in control conditions. Cluster A and B comprise 137 and 120 genes, respectively. The continuous black line shows the mean of mRNA expression in different conditions. (a’, b’) Gene ontology analysis of biological processes for genes of clusters A and B. (C, D) Heatmap of mRNAs expressed differentially in different conditions for different classes of coded protein. (\**p* < 0.05, \*\**p* < *0.01*, *\*\*\*p* < 0.001). Color intensity shows the Z-score for differential expression.

We found a significant difference in the group of genes that were reduced in PUR cultures and restored in OCM cultures. For one group of these genes, expression in GABAergic and pyramidal neurons in CTRL conditions were comparable (Fig. 8A; n=137). Another second group of genes differed in that expression in GABAergic neurons was systematically higher than in pyramidal cells in CTRL cultures (Fig. 8B; n=120). GO term analysis of the identities of the first group of genes (Fig. 8a’) linked them to processes including negative regulation of protein kinase activity and apoptosis signaling pathways. In contrast, the identities of the second group of genes (Fig. 8b’) evoked biological processes including the membrane transport of zinc, K^+^ channels, regulation of membrane potential and action potential and G-protein coupled receptor signaling pathways.

Finally we present target genes whose expression was affected by the absence of glial cells in PUR cultures and inversely by OCM and which are linked to GABAergic cell physiology. These target transcripts code for ion channels, transporters, synaptic markers, vesicle trafficking, cytoskeleton remodeling, cell adhesion molecules, growth factors and signaling (Fig. 8C) and were mostly up-regulated in CTRL and OCM neurons. They included Na^+^ channels, *Scn1a* (Nav1.1) and *Scn1b* (ß1Nav) and K^+^ channels, *Kcna1* (K*v*1.1), *Kcnab1* (ß1Kv), *Kcnk3* (Task-1), *Kcnip1* (KChip-1, a K^+^ channel modulatory protein). Genes encoding zinc transporters (*Slc39a11, Slc30a4* and *Slc30a3*), were upregulated by OCM, while a Na^+^/K^+^/Ca^2+^ exchanger (*Slc24a2*) was one of the few transcripts downregulated by OCM. Among genes coding for signaling molecules, growth factors and receptors, the kinases *Nek7, Pak1* and *Akt1*, and growth factors *Vegfb, Pdfga, Fgf9 and Rara* were up-regulated in OCM culture conditions. Only the protein kinase C binding protein *Nell2* and the neuropeptide hormone Proenkephalin (*Penk*) were downregulated by OCM.

## DISCUSSION

Data presented here suggests that factors released by oligodendrocytes modulate the transcriptome, electrical phenotype and morphology of GABAergic neurons. The absence of glial cells from purified neuron cultures reduced synaptic activity and action potential firing of GABAergic neurons and many genes were downregulated. Processes linked to these changes included synapse assembly, action potential generation and transmembrane ion transport. Our results should help identify some of the molecular targets by which oligodendrocytes modulate GABAergic neuron excitability and synaptic function.

### A hippocampal culture model permits investigation of regulation of interneuron morphology, excitability and firing properties by glial factors

This study was based on dissociated hippocampal cell cultures. Recordings were made from (i) glutamatergic and GABAergic neurons in the presence of glial cells (CTRL), (ii) GABAergic neurons in the absence of glial cells (PUR) and (iii), GABAergic neurons in cultures without glial cells but with added oligodendroglial conditioned medium (OCM). Our data showed (Fig. 2) GABAergic neurons in culture conserved properties, including a relatively depolarized resting potential, and action potentials of short duration followed by a prominent AHP, which distinguish them from hippocampal pyramidal cells (Spruston and Johnston 1992; Fricker et al. 1999; Staff et al. 2000; Hu et al. 2014).

The frequency of excitatory synaptic events impinging on hippocampal GABAergic neurons was strongly reduced in the absence of glial cells. This is consistent with data showing that glutamatergic synaptic transmission is enhanced by factors secreted by glia (Turko et al. 2019) including astrocytes (Baldwin and Eroglu 2017). Here we demonstrate for the first time that oligodendrocytes regulate synaptic excitation of GABAergic interneurons. Adding OCM to PUR cultures partly restored EPSC frequencies. The absence of astrocyte related factors from the conditioned medium may appear to reduce the strength of the oligodendrocyte related effect.

Anatomical analysis revealed that oligodendrocyte secreted factors (OCM) enhanced the complexity of dendritic arbors, although dendritic length differed little between cultures that did (CTRL) or did not (PUR) contain glial cells. Novel synapses formed on more complex dendrites in OCM cultures may contribute to the increased EPSC frequency in these conditions.

We found similar electrophysiological properties for GABAergic neurons grown in CTRL and OCM cultures, notably a low input resistance and high rheobase. Rheobase was reduced and AP firing frequency significantly depressed in PUR cultures. The phenotype of low intrinsic excitability corresponds to mature neurons expressing a large array of voltage dependent or independent ion channels. Glial factors may operate homeostatically so that neurons adapt to levels of spontaneous firing in cultures with dense synaptic connections.

We did not attempt to identify factors released by oligodendrocytes, and their precursor cells, which mediated the effects of OCM on GABAergic neurons. Candidate molecules may include the proteoglycan NG2 (Sakry et al. 2014), FGF2 (Birey et al. 2015), or BDNF (Jang et al. 2019) which modulate glutamatergic neurotransmission on pyramidal cells but their effects on GABAergic neurons are not known. Our mass spectrometry analysis has shown that contactin-1, together with RPTPß/phosphacan or tenascin R, is present in a previous OCM, and influences early formation of Na+ channel clusters on axons of GABAergic cells (Dubessy et al. 2019). Cell adhesion molecules and extracellular matrix proteins secreted by oligodendrocytes form peri-nodal complexes that transmit signals to neurons which could influence their physiology and connectivity (Fawcett, Oohashi and Pizzorusso, 2019).

### Technical points

This work combined patch clamp recordings with single neuron RNA-seq analysis. Relatively few studies have demonstrated strong correlations between single-neuron transcriptomic profiles and electrophysiological phenotypes (Cadwell et al. 2016; Földy et al. 2016; Fuzik et al. 2016; Muñoz-Manchado et al. 2018; Scala et al. 2019). We took several steps to ensure and verify the validity of our results. The duration of patch electrode recordings was deliberately limited to ~10 min in order to minimize perturbation of the transcriptome (Fuzik et al. 2016). When possible patch electrodes targeted isolated neuronal somata to reduce possible mRNA contamination from adjacent cells (Tripathy et al. 2017).

As for previously published single-cell RNAseq datasets, the number of sequenced reads per cell was found to be positively correlated with detected transcript counts and did not reach a plateau (Cadwell et al. 2016; Tasic et al. 2016; Tripathy et al. 2017). We attempted to validate our data by cross-correlating transcripts detected for recorded pyramidal and inhibitory neurons with expected profiles for these cell types. Most GABAergic neurons expressed molecular markers, including peptides and Ca-binding proteins, specific to known subclasses of these cells (Zeisel et al. 2015; Gouwens et al. 2020). Larger numbers of sequenced cells would have permitted enhanced statistics to assure data quality even if links between specific transcripts and identified cell types tends to support our approach.

Single neuron transcriptomes obtained in this way helped us define a global view of processes initiated by oligodendrocyte conditioned medium. They showed glial factors modify the transcriptome of GABAergic neurons to change intrinsic electrophysiological properties, AP generation, EPSC frequencies and dendritic anatomy.

### Gene expression in hippocampal neuron types

Transcriptomic data are defining the classification of cortical neurons (Zeisel et al. 2015; Cembrowski et al. 2016; Harris et al. 2018; Sugino et al. 2019; Yuste et al. 2020). A recent study based on Patch-seq data from several 1000s of cells may offer the best current correspondence between transcriptomic, anatomical and electrophysiological data for GABAergic mouse cortical neurons (Gouwens et al. 2020). Our data can be interpreted in the light of those studies. Most GABAergic neurons studied in the hippocampal cultures studied here expressed Somatostatin (SST) associated with other peptide markers. Several subtypes of hippocampal inhibitory cells express SST including long-range inhibitory neurons which possess myelinated axons (and also express *Calbindin* and *Npy*), oriens-lacunosum moleculare (O-LM) interneurons (also *Elfn1* and *Pnoc*), or oriens-bistratified neurons (also *Tac1*, *Npy*, *Satb1* and *Erbb4*) (Somogyi and Klausberger 2005; Jinno 2009; Harris et al. 2018). We found some cells expressed genes for *Sst*, *Pnoc* and *Pvalb* (Jinno and Kosaka 2000; Jinno 2009; Harris et al. 2018). A minority of GABAergic neurons expressed genes for *Reelin* and *NPY* as do neurogliaform cells (Pelkey et al., 2017).

GABAergic neuron data revealed genes coding for proteins relevant to specific aspects of inhibitory cell physiology. They included Nav1.1 (*Scn1a*), Kv3.2 (*Kcnc2*) and Task-1(*Kcnk3*) in neurons with PV and/or SST genes (Chow et al. 1999; Torborg et al. 2006; Lorincz and Nusser 2008). Kv3 channels with fast kinetics curtail action potentials permitting sustained firing at high frequencies (Rudy and McBain 2001; Gu et al. 2018; Hu et al. 2018). Task-1 forms K^+^ permeable leak channels which contribute to resting potential and membrane resistance (Okaty et al. 2009). We found high levels of genes for the zinc transporters, ZnT3 (*Slc30a3*) and ZnT4 (*Slc30a4*), which are found in SST-containing interneurons (Paul et al. 2017). ZnT3 is a vesicular transporter which may contribute to the co-release of zinc in synaptic vesicles with GABA (McAllister and Dyck 2017).

### Correlation between gene expression and electrophysiological parameters

We attempted to link expression of genes for ion channels and neurotransmitter receptors with elements of electrophysiological phenotypes using a correlation-based analysis on data from different cell types and culture conditions. The analysis suggests genes for α1, ß1 and α3 subunits of the Na^+^, K^+^ -ATPase, *Atp1a1*, *Atp1b1* and *Atp1a3*, are linked to neuronal input resistance. The Na^+^/K^+^-ATPase maintains transmembrane ionic gradients and resting membrane thus affecting neuronal excitability (Larsen et al. 2016). The Na-channel subunit *Scn2a* (Nav1.2) was found to be correlated with action potential width, neuronal input resistance, and the membrane time constant, tau. A cluster including several K^+^ channels (*Kcnc2*, *Kcnk3* and *Kcnip-1*) and two Zinc transporters (*Slc30a3 and Slc30a*), was expressed selectively in GABAergic neurons, and correlated with AP threshold, sag ratio and the after-hyperpolarization amplitude. We should note that these correlations do not imply causality and caution that class-driven correlations are an important confound in our dataset. Some within-cell-type correlations may have been missed (Bomkamp et al. 2019).

### Biological processes affected by glial cells and oligodendroglial secreted factors

Clustering genes with similar patterns of altered expression revealed GO process terms regulated by factors in OCM. Processes identified in this way matched quite efficiently with changes in GABAergic neuron phenotype inferred from electrophysiological and anatomical observations. Enriched processes included synapse assembly, action potential generation, trans-membrane transport of ions specifically zinc, and kinase signaling. They derived from differential expression of K^+^ channel genes, including *Kcna1* (Kv1.1), *Kcnab1* (Kv ß1chain), *Kcnip1* (KChIP) and *Kcnk3* (Task-1), and Na^+^ channel genes, including *Scn1a* (Nav1.1) and *Scn1b* (Nav ß1chain). We found two kinases which were upregulated in OCM. Nek7 is involved in microtubule polymerization during the formation of PV+ interneuron connections (Hinojosa et al. 2018). Akt1, mediates the effects of growth factors to regulate neuronal survival, actin polymerization, synaptic transmission and to initiate myelination by oligodendrocytes (Wang et al. 2003; Lai et al. 2006; Goebbels et al. 2017).

In conclusion, our study provides new insights into communication between glial cells and neurons showing that factors secreted by oligodendrocytes induce transcriptomic changes which may modulate the physiology and anatomy of GABAergic neurons. Further work, possibly including single-cell transcriptomics from oligodendrocytes, should focus on the identity of secreted factors. Studies extended to characterize signaling from oligodendrocytes and their precursors to pyramidal cells could profitably establish general principles underlying the postnatal myelination of cortical axons. Understanding these principles may help design therapies to enhance myelin repair capacity in demyelinating diseases such as multiple sclerosis.

## Acknowledgements

We thank Marie-Stéphane Aigrot and Loane Wallon, Claire Lovo from ICM Quant, Yannick Marie and Emeline Mundwiller from iGenSeq, and François-Xavier Lejeune from iCONICS, for technical support. We thank Bernard Zalc, Richard Miles and Anne Desmazieres for discussion and critical reading of the manuscript. This work was supported by the French MS research foundation ARSEP (Aide à la Recherche sur la Sclérose en Plaques), Bouvet-Labruyère prize to NSF, and Biogen funding to EM.

## REFERENCES

Arancibia-Cárcamo IL, Ford MC, Cossell L, Ishida K, Tohyama K, Attwell D. 2017. Node of Ranvier length as a potential regulator of myelinated axon conduction speed. Elife. 6.

Baldwin KT, Eroglu C. 2017. Molecular mechanisms of astrocyte-induced synaptogenesis. Curr Opin Neurobiol. 45:113–120.

Bates D, Mächler M, Bolker B, Walker S. 2015. Fitting Linear Mixed-Effects Models Using lme4. Journal of Statistical Software 67, 1–48.

Barres BA, Raff MC. 1993. Proliferation of oligodendrocyte precursor cells depends on electrical activity in axons. Nature. 361(6409):258–260.

Battefeld A, Klooster J, Kole MHP. 2016. Myelinating satellite oligodendrocytes are integrated in a glial syncytium constraining neuronal high-frequency activity. Nat Commun. 7:11298.

Bechler ME, Swire M, Ffrench-Constant C. 2018. Intrinsic and adaptive myelination-A sequential mechanism for smart wiring in the brain. Dev Neurobiol. 78:68–79.

Birey F, Kloc M, Chavali M, Hussein I, Wilson M, Christoffel DJ, Chen T, Frohman MA, Robinson JK, Russo SJ, Maffei A, Aguirre A. 2015. Genetic and Stress-Induced Loss of NG2 Glia Triggers Emergence of Depressive-like Behaviors through Reduced Secretion of FGF2. Neuron. 88:941–956.

Bomkamp C, Tripathy SJ, Bengtsson Gonzales C, Hjerling-Leffler J, Craig AM, Pavlidis P. 2019. Transcriptomic correlates of electrophysiological and morphological diversity within and across excitatory and inhibitory neuron classes. PLoS Comput Biol. 15:e1007113.

Bonetto G, Hivert B, Goutebroze L, Karagogeos D, Crépel V, Faivre-Sarrailh C. 2019. Selective Axonal Expression of the Kv1 Channel Complex in Pre-myelinated GABAergic Hippocampal Neurons. Front Cell Neurosci. 13:222.

Cadwell CR, Palasantza A, Jiang X, Berens P, Deng Q, Yilmaz M, Reimer J, Shen S, Bethge M, Tolias KF, Sandberg R, Tolias AS. 2016. Electrophysiological, transcriptomic and morphologic profiling of single neurons using Patch-seq. Nat Biotechnol. 34:199–203.

Cembrowski MS, Wang L, Sugino K, Shields BC, Spruston N. 2016. Hipposeq: a comprehensive RNA-seq database of gene expression in hippocampal principal neurons. Elife. 5:e14997.

Chow A, Erisir A, Farb C, Nadal MS, Ozaita A, Lau D, Welker E, Rudy B. 1999. K(+) channel expression distinguishes subpopulations of parvalbumin- and somatostatin-containing neocortical interneurons. J Neurosci. 19:9332–9345.

Demerens C, Stankoff B, Logak M, Anglade P, Allinquant B, Couraud F, Zalc B, Lubetzki C. 1996. Induction of myelination in the central nervous system by electrical activity. Proc Natl Acad Sci USA. 93(18):9887–9892.

Dobin A, Davis CA, Schlesinger F, Drenkow J, Zaleski C, Jha S, Batut P, Chaisson M, Gingeras TR. 2013. STAR: ultrafast universal RNA-seq aligner. Bioinformatics. 29:15–21.

Dubessy A-L, Mazuir E, Rappeneau Q, Ou S, Abi Ghanem C, Piquand K, Aigrot M-S, Thétiot M, Desmazières A, Chan E, Fitzgibbon M, Fleming M, Krauss R, Zalc B, Ranscht B, Lubetzki C, Sol-Foulon N. 2019. Role of a Contactin multi-molecular complex secreted by oligodendrocytes in nodal protein clustering in the CNS. Glia. 67:2248–2263.

Fawcett JW, Oohashi T and Pizzorusso T 2019. The roles of perineuronal nets and the perinodal extracellular matrix in neuronal function. Nat Rev Neurosci. 20: 451–465.

Földy C, Darmanis S, Aoto J, Malenka RC, Quake SR, Südhof TC. 2016. Single-cell RNAseq reveals cell adhesion molecule profiles in electrophysiologically defined neurons. Proc Natl Acad Sci USA. 113:E5222–5231.

Freeman SA, Desmazières A, Fricker D, Lubetzki C, Sol-Foulon N. 2016. Mechanisms of sodium channel clustering and its influence on axonal impulse conduction. Cell Mol Life Sci. 73:723–735.

Freeman SA, Desmazières A, Simonnet J, Gatta M, Pfeiffer F, Aigrot MS, Rappeneau Q, Guerreiro S, Michel PP, Yanagawa Y, Barbin G, Brophy PJ, Fricker D, Lubetzki C, Sol-Foulon N. 2015. Acceleration of conduction velocity linked to clustering of nodal components precedes myelination. Proc Natl Acad Sci USA. 112:E321–328.

Fricker D, Verheugen JA, Miles R. 1999. Cell-attached measurements of the firing threshold of rat hippocampal neurones. J Physiol. 517:791–804.

Frühbeis C, Fröhlich D, Kuo WP, Amphornrat J, Thilemann S, Saab AS, Kirchhoff F, Möbius W, Goebbels S, Nave K-A, Schneider A, Simons M, Klugmann M, Trotter J, Krämer-Albers EM. 2013. Neurotransmitter-triggered transfer of exosomes mediates oligodendrocyte-neuron communication. PLoS Biol. 11:e1001604.

Fünfschilling U, Supplie LM, Mahad D, Boretius S, Saab AS, Edgar J, Brinkmann BG, Kassmann CM, Tzvetanova ID, Möbius W, Diaz F, Meijer D, Suter U, Hamprecht B, Sereda MW, Moraes CT, Frahm J, Goebbels S, Nave KA. 2012. Glycolytic oligodendrocytes maintain myelin and long-term axonal integrity. Nature. 485:517–521.

Fuzik J, Zeisel A, Máté Z, Calvigioni D, Yanagawa Y, Szabó G, Linnarsson S, Harkany T. 2016. Integration of electrophysiological recordings with single-cell RNA-seq data identifies neuronal subtypes. Nat Biotechnol. 34:175–183.

Goebbels S, Wieser GL, Pieper A, Spitzer S, Weege B, Yan K, Edgar JM, Yagensky O, Wichert SP, Agarwal A, Karram K, Renier N, Tessier-Lavigne M, Rossner MJ, Káradóttir RT, Nave KA. 2017. A neuronal PI(3,4,5)P3-dependent program of oligodendrocyte precursor recruitment and myelination. Nat Neurosci. 20:10–15.

Gouwens NW, Sorensen SA, Baftizadeh F, Budzillo A, Lee BR, Jarsky T, Alfiler L, Arkhipov A, Baker K, Barkan E, et al. 2020. Toward an integrated classification of neuronal cell types: morphoelectric and transcriptomic characterization of individual GABAergic cortical neurons. bioRxiv. 2020.02.03.932244.

Gu Y, Servello D, Han Z, Lalchandani RR, Ding JB, Huang K, Gu C. 2018. Balanced Activity between Kv3 and Nav Channels Determines Fast-Spiking in Mammalian Central Neurons. iScience. 9:120–137.

Hu H, Gan J, Jonas P 2014. Interneurons. Fast-spiking, parvalbumin^+^ GABAergic interneurons: from cellular design to microcircuit function. Science. 345: 1255263.

Harris KD, Hochgerner H, Skene NG, Magno L, Katona L, Bengtsson Gonzales C, Somogyi P, Kessaris N, Linnarsson S, Hjerling-Leffler J. 2018. Classes and continua of hippocampal CA1 inhibitory neurons revealed by single-cell transcriptomics. PLoS Biol. 16:e2006387.

Hinojosa AJ, Deogracias R, Rico B. 2018. The Microtubule Regulator NEK7 Coordinates the Wiring of Cortical Parvalbumin Interneurons. Cell Rep. 24:1231–1242.

Hu H, Roth FC, Vandael D, Jonas P. 2018. Complementary Tuning of Na+ and K+ Channel Gating Underlies Fast and Energy-Efficient Action Potentials in GABAergic Interneuron Axons. Neuron. 98:156–165.e6.

Huang DW, Sherman BT, Tan Q, Collins JR, Alvord WG, Roayaei J, Stephens R, Baseler MW, Lane HC, Lempicki RA. 2007. The DAVID Gene Functional Classification Tool: a novel biological module-centric algorithm to functionally analyze large gene lists. Genome Biol. 8: R183.

Huang L-W, Simonnet J, Nassar M, Richevaux L, Lofredi R, Fricker D. 2017. Laminar Localization and Projection-Specific Properties of Presubicular Neurons Targeting the Lateral Mammillary Nucleus, Thalamus, or Medial Entorhinal Cortex. eNeuro. 4:0370–16.

Jang M, Gould E, Xu J, Kim EJ, Kim JH. 2019. Oligodendrocytes regulate presynaptic properties and neurotransmission through BDNF signaling in the mouse brainstem. Elife. 8:e42156.

Jinno S. 2009. Structural organization of long-range GABAergic projection system of the hippocampus. Front Neuroanat. 3:13.

Jinno S, Kosaka T. 2000. Colocalization of parvalbumin and somatostatin-like immunoreactivity in the mouse hippocampus: Quantitative analysis with optical disector. The J Comp Neurol. 428:377–388.

Kaplan MR, Meyer-Franke A, Lambert S, Bennett V, Duncan ID, Levinson SR, Barres BA. 1997. Induction of sodium channel clustering by oligodendrocytes. Nature. 386:724–728.

Lai W-S, Xu B, Westphal KGC, Paterlini M, Olivier B, Pavlidis P, Karayiorgou M, Gogos JA. 2006. Akt1 deficiency affects neuronal morphology and predisposes to abnormalities in pre-frontal cortex functioning. Proc Natl Acad Sci USA. 103:16906–16911.

Larsen BR, Stoica A, MacAulay N. 2016. Managing Brain Extracellular K(+) during Neuronal Activity: The Physiological Role of the Na(+)/K(+)-ATPase Subunit Isoforms. Front Physiol. 7:141.

Lee Y, Morrison BM, Li Y, Lengacher S, Farah MH, Hoffman PN, Liu Y, Tsingalia A, Jin L, Zhang P-W, Pellerin L, Magistretti PJ, Rothstein JD. 2012. Oligodendroglia metabolically support axons and contribute to neurodegeneration. Nature. 487:443–448.

Lorincz A, Nusser Z. 2008. Cell-type-dependent molecular composition of the axon initial segment. J Neurosci. 28:14329–14340.

Love MI, Huber W, Anders S. 2014. Moderated estimation of fold change and dispersion for RNA-seq data with DESeq2. Genome Biol. 15:550.

Mazuir E, Dubessy A-L, Wallon L, Aigrot M-S, Lubetzki C, Sol-Foulon N. 2020. Generation of Oligodendrocytes and Oligodendrocyte-Conditioned Medium for Co-Culture Experiments. J Vis Exp. (156).

McAllister BB, Dyck RH. 2017. Zinc transporter 3 (ZnT3) and vesicular zinc in central nervous system function. Neurosci Biobehav Rev. 80:329–350.

McCarthy KD, de Vellis J. 1980. Preparation of separate astroglial and oligodendroglial cell cultures from rat cerebral tissue. J Cell Biol. 85:890–902.

McKenzie IA, Ohayon D, Li H, de Faria JP, Emery B, Tohyama K, Richardson WD. 2014. Motor skill learning requires active central myelination. Science. 346:318–322.

Monje M. 2018. Myelin Plasticity and Nervous System Function. Annu Rev Neurosci. 41:61–76.

Muñoz-Manchado AB, Bengtsson Gonzales C, Zeisel A, Munguba H, Bekkouche B, Skene NG, Lönnerberg P, Ryge J, Harris KD, Linnarsson S, Leffler JH. 2018. Diversity of Interneurons in the Dorsal Striatum Revealed by Single-Cell RNA Sequencing and PatchSeq. Cell Rep. 24:2179–2190.e7.

Noli L, Capalbo A, Ogilvie C, Khalaf Y, Ilic D. 2015. Discordant Growth of Monozygotic Twins Starts at the Blastocyst Stage: A Case Study. Stem Cell Reports. 5:946–953.

Okaty BW, Miller MN, Sugino K, Hempel CM, Nelson SB. 2009. Transcriptional and electrophysiological maturation of neocortical fast-spiking GABAergic interneurons. J Neurosci. 29:7040–7052.

Paul A, Crow M, Raudales R, He M, Gillis J, Huang ZJ. 2017. Transcriptional Architecture of Synaptic Communication Delineates GABAergic Neuron Identity. Cell. 171:522–539.e20.

Pelkey KA, Chittajallu R, Craig MT, Tricoire L, Wester JC, McBain CJ. 2017. Hippocampal GABAergic Inhibitory Interneurons. Physiol Rev. 97:1619–1747.

Qiu S, Luo S, Evgrafov O, Li R, Schroth GP, Levitt P, Knowles JA, Wang K. 2012. Single-neuron RNA-Seq: technical feasibility and reproducibility. Front Genet. 3:124.

Rudy B, McBain CJ. 2001. Kv3 channels: voltage-gated K+ channels designed for high-frequency repetitive firing. Trends Neurosci. 24:517–526.

Saab AS, Tzvetavona ID, Trevisiol A, Baltan S, Dibaj P, Kusch K, Möbius W, Goetze B, Jahn HM, Huang W, Steffens H, Schomburg ED, Pérez-Samartín A, Pérez-Cerdá F, Bakhtiari D, Matute C, Löwel S, Griesinger C, Hirrlinger J, Kirchhoff F, Nave KA. 2016. Oligodendroglial NMDA Receptors Regulate Glucose Import and Axonal Energy Metabolism. Neuron. 91:119–132.

Sakry D, Yigit H, Dimou L, Trotter J. 2015. Oligodendrocyte precursor cells synthesize neuro-modulatory factors. PLoS ONE. 10:e0127222.

Scala F, Kobak D, Shan S, Bernaerts Y, Laturnus S, Cadwell CR, Hartmanis L, Froudarakis E, Castro JR, Tan ZH, Papadopoulos S, Patel SS, Sandberg R, Berens P, Jiang X, Tolias AS. 2019. Layer 4 of mouse neocortex differs in cell types and circuit organization between sensory areas. Nat Commun. 10:4174.

Seidl AH. 2014. Regulation of conduction time along axons. Neuroscience. 276:126–134.

Sherman DL, Brophy PJ. 2005. Mechanisms of axon ensheathment and myelin growth. Nat Rev Neurosci. 6:683–690.

Somogyi P, Klausberger T. 2005. Defined types of cortical interneurone structure space and spike timing in the hippocampus. J Physiol. 562:9–26.

Spruston N, Johnston D. 1992. Perforated patch-clamp analysis of the passive membrane properties of three classes of hippocampal neurons. J Neurophysiol. 67:508–529.

Staff NP, Jung HY, Thiagarajan T, Yao M, Spruston N. 2000. Resting and active properties of pyramidal neurons in subiculum and CA1 of rat hippocampus. J Neurophysiol. 84: 2398–2408.

Stedehouder J, Brizee D, Shpak G, Kushner SA. 2018. Activity-Dependent Myelination of Parvalbumin Interneurons Mediated by Axonal Morphological Plasticity. J Neurosci. 38:3631–3642.

Sugino K, Clark E, Schulmann A, Shima Y, Wang L, Hunt DL, Hooks BM, Tränkner D, Chan-drashekar J, Picard S, Lemire AL, Spruston N, Hantman AW Nelson SB. 2019. Mapping the transcriptional diversity of genetically and anatomically defined cell populations in the mouse brain. Elife. 8:e38619.

Tasic B, Menon V, Nguyen TN, Kim TK, Jarsky T, Yao Z, Levi B, Gray LT, Sorensen SA, Dolbeare T, Bertagnolli D, Goldy J, Shapovalova N, Parry S, Lee C, Smith K, Bernard A, Madisen L, Sunkin SM, Hawrylycz M, Koch C, Zeng H. 2016. Adult mouse cortical cell taxonomy revealed by single cell transcriptomics. Nat Neurosci. 19:335–346.

Torborg CL, Berg AP, Jeffries BW, Bayliss DA, McBain CJ. 2006. TASK-like conductances are present within hippocampal CA1 stratum oriens interneuron subpopulations. J Neurosci. 26:7362–7367.

Tripathy SJ, Toker L, Li B, Crichlow C-L, Tebaykin D, Mancarci BO, Pavlidis P. 2017. Transcriptomic correlates of neuron electrophysiological diversity. PLoS Comput Biol. 13:e1005814.

Turko P, Groberman K, Browa F, Cobb S, Vida I. 2019. Differential Dependence of GABAergic and Glutamatergic Neurons on Glia for the Establishment of Synaptic Transmission. Cereb Cortex. 29:1230–1243.

Uematsu M, Hirai Y, Karube F, Ebihara S, Kato M, Abe K, Obata K, Yoshida S, Hirabayashi M, Yanagawa Y, Kawaguchi Y. 2008. Quantitative chemical composition of cortical GABAergic neurons revealed in transgenic venus-expressing rats. Cereb Cortex. 18(2):315–330.

Wang Q, Liu L, Pei L, Ju W, Ahmadian G, Lu J, Wang Y, Liu F, Wang YT. 2003. Control of synaptic strength, a novel function of Akt. Neuron. 38:915–928.

Wickham H. 2016. Ggplot2: Elegant Graphics for Data Analysis. Springer-Verlag New York. ISBN 978-3-319-24277-4

Wilson MD, Sethi S, Lein PJ, Keil KP. 2017. Valid statistical approaches for analyzing sholl data: Mixed effects versus simple linear models. J Neurosci Methods. 279:33–43.

Xin W, Mironova YA, Shen H, Marino RAM, Waisman A, Lamers WH, Bergles DE, Bonci A. 2019. Oligodendrocytes Support Neuronal Glutamatergic Transmission via Expression of Glutamine Synthetase. Cell Rep. 27:2262–2271.e5.

Yuste R, Hawrylycz M, Aalling N, Aguilar-Valles A, Arendt D, Arnedillo RA, Ascoli GA, Bielza C, Bokharaie V, Bergmann TB, et al. 2020. A community-based transcriptomics classification and nomenclature of neocortical cell types. Nat Neurosci. PMID: 32839617.

Zeisel A, Muñoz-Manchado AB, Codeluppi S, Lönnerberg P, La Manno G, Juréus A, Marques S, Munguba H, He L, Betsholtz C, Rolny C, Castelo-Branco G, Hjerling-Leffler J, Linnarsson S. 2015. Brain structure. Cell types in the mouse cortex and hippocampus revealed by single-cell RNA-seq. Science. 347:1138–1142.

