## Supplementary figures and images for "Oligodendrocyte secreted factors shape hippocampal GABAergic neuron transcriptome and physiology"

### Supplemental Figure 1

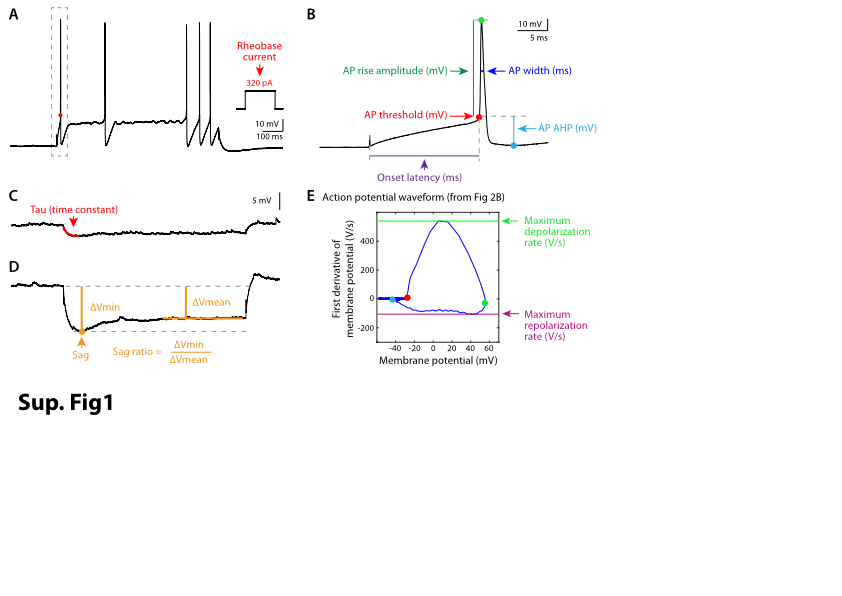

### Supplemental Figure 2

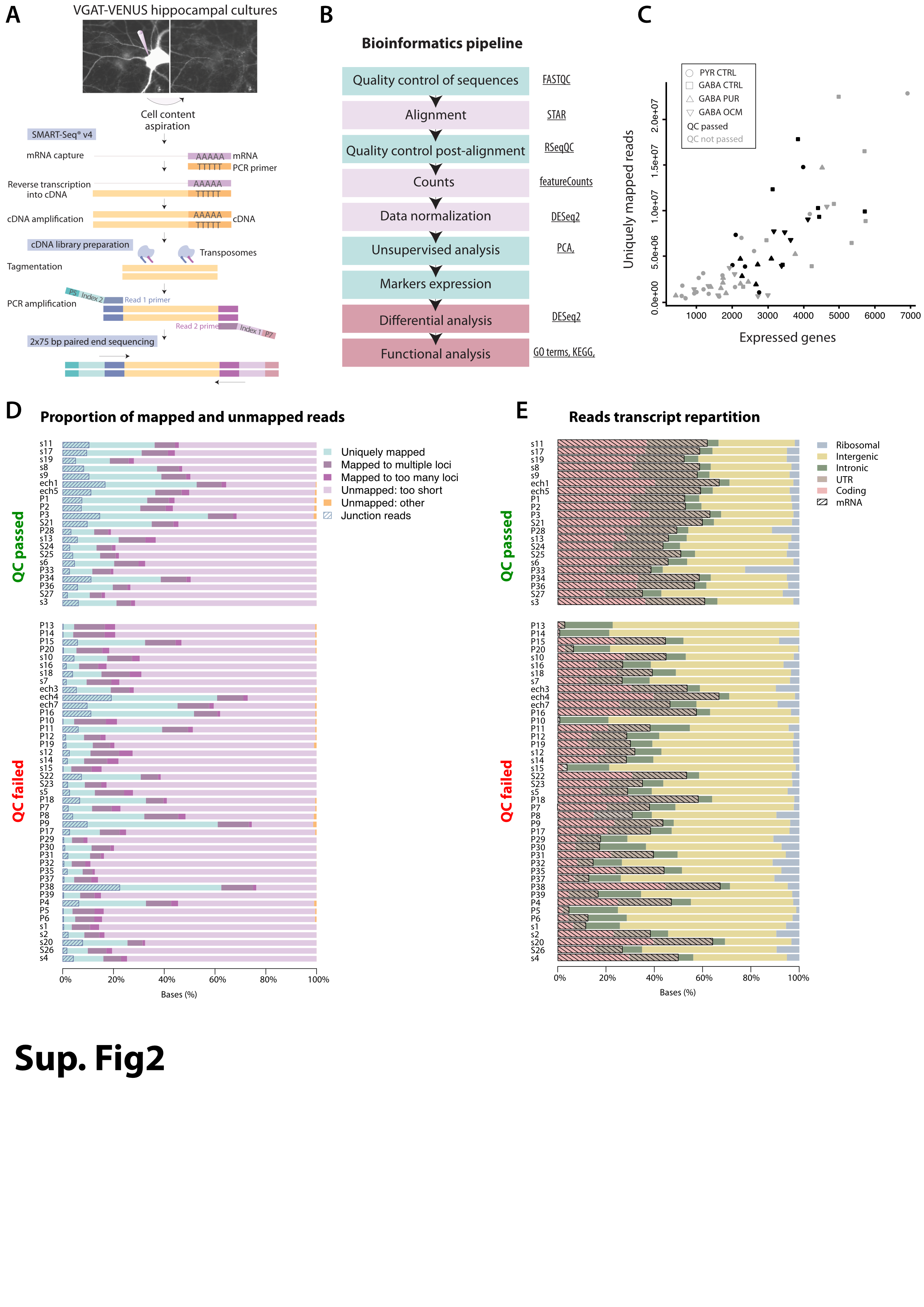

### Supplemental Figure 3

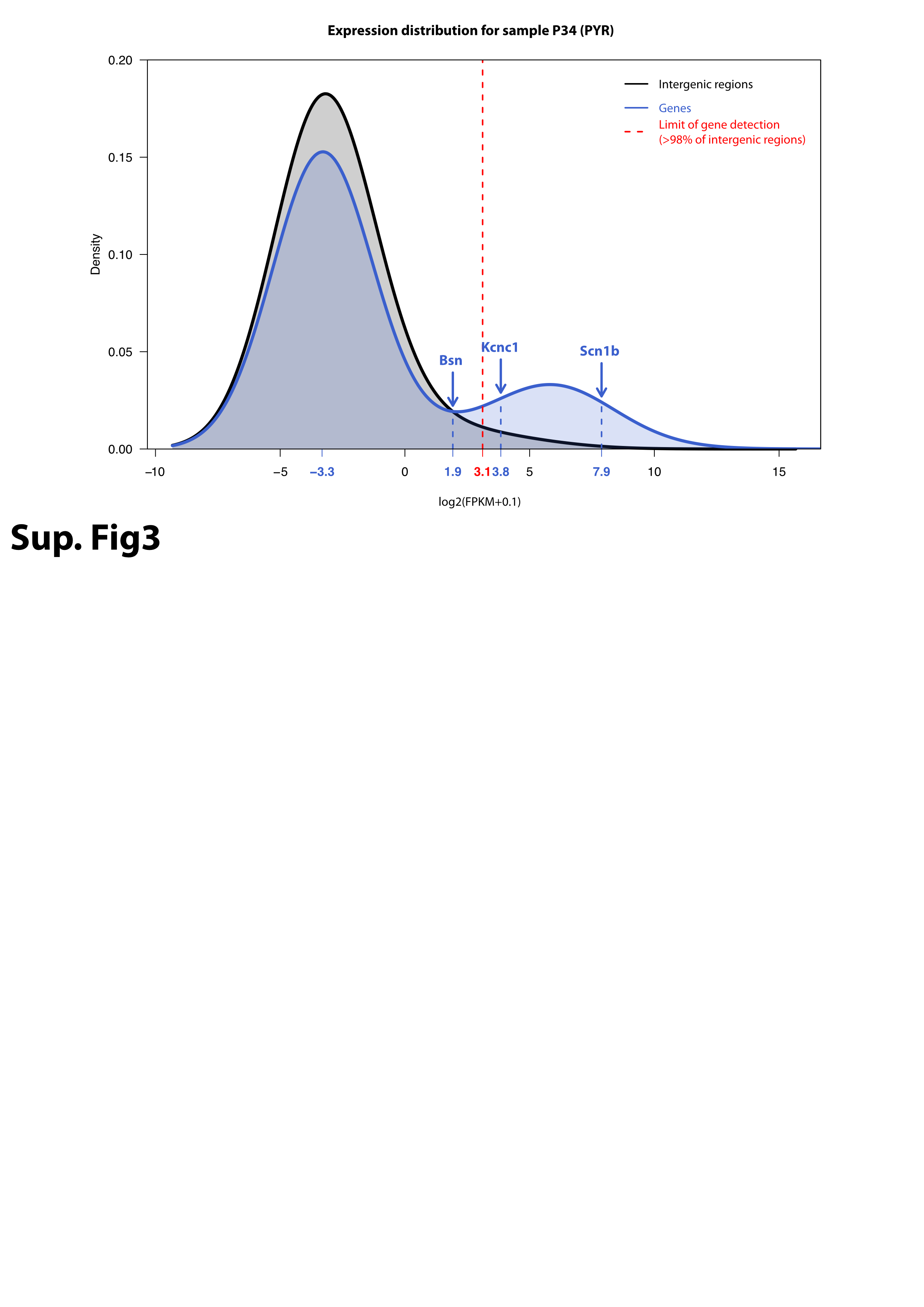

### Supplemental Figure 4

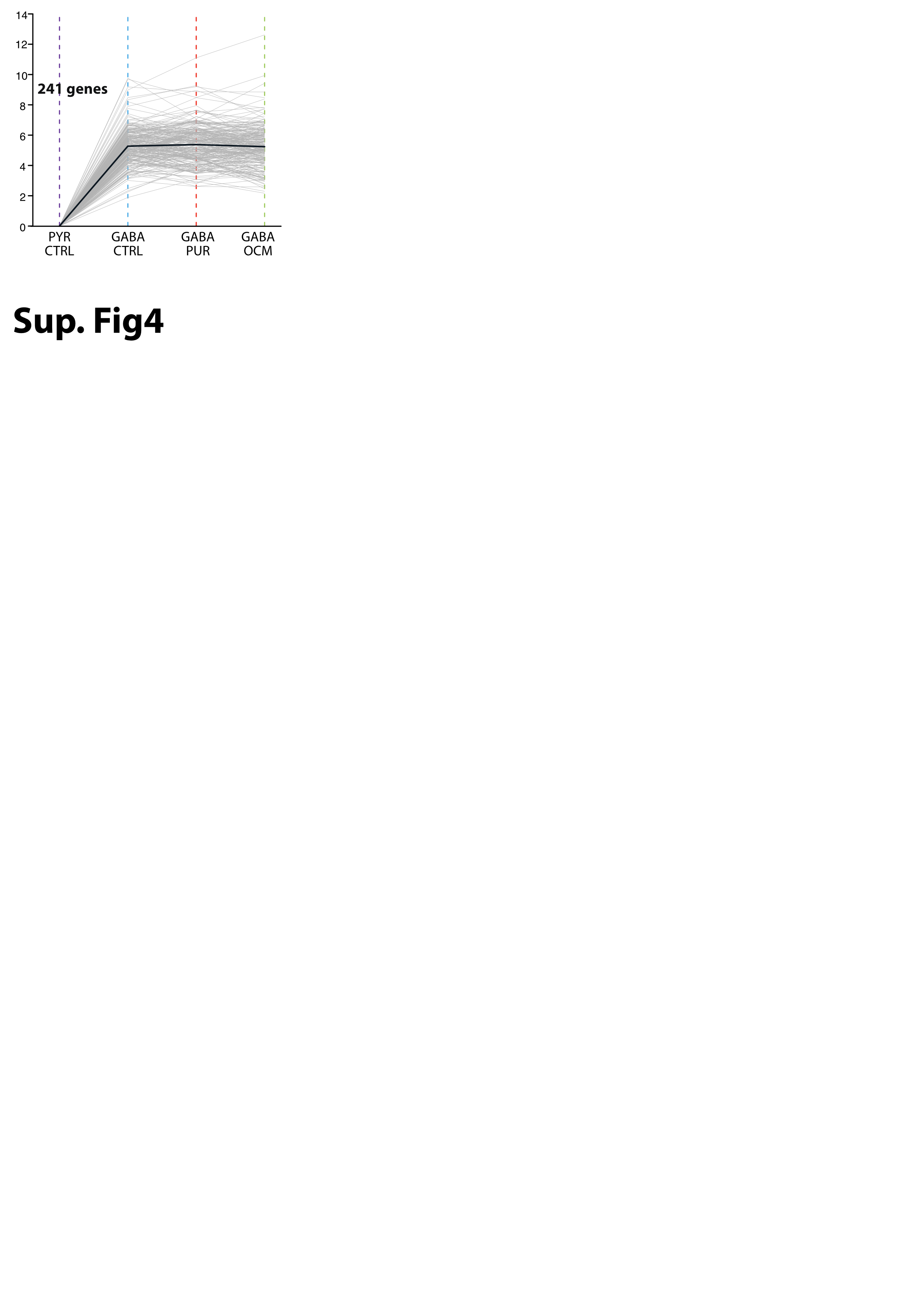
