## supplementary legend for "Oligodendrocyte secreted factors shape hippocampal GABAergic neuron transcriptome and physiology"

**Supplementary material**

**Sup. Figure 1: (A)** Voltage response of an inhibitory cell to rheobase injected current, the minimal intensity needed to initiate an action potential (AP). In this example, rheobase is 320 pA. The AP voltage threshold is defined as the point at the foot of the first AP where dV/dt exceeds 30 mV/ms, indicated by a red dot. **(B)** The first AP at rheobase (dashed grey area from **A)**. The following measures of active neuronal properties presented in **Fig. 1** and **2** are derived from analysis of this waveform: Onset latency (purple line), AP threshold (red dot), AP rise amplitude (green line), AP width (dark blue line), AP afterhyperpolarization (AHP, light blue dot and line). Green point represents AP peak. **(C)** The membrane time constant (Tau) is the shorter time constant determined by fitting a double exponential function to a membrane response (-10 mV frombaseline) to a small hyperpolarizing current. **(D)** Sag ratio is related to the I_h_ current. It is the average of three measures of ΔVmin/ΔVmean around -100 mV (ΔVmin and ΔVmean are relative to baseline). **(E)** Phase plot representation of the first action potential at rheobase, as the first derivative of membrane potential against membrane potential. Dots indicate membrane potential parameters as in **(B):** the derivative increases from the threshold (red dot) to maximum depolarization rate (maximum slope of AP depolarization, green line) and then decreases to the AP peak (green dot). After peak, the derivative decreases to reach the maximum repolarization rate (maximum slope of AP repolarization) and then further to AP AHP (blue dot), the most negative voltage point of the AP waveform.

**Sup. Figure 2: (A)** Overview of transcript library preparation. Cytosol content was extracted from hippocampal neurons by a patch pipette. SMART-Seq v4 technology was used for mRNA capture, reverse transcription and cDNA amplification. Full-length cDNA was processed with the Nextera XT DNA Library Preparation Kit from Illumina to generate multiplex sequencing libraries. Next-generation sequencing was used to produce libraries with the Illumina NextSeq500 after a 2x75bp paired end sequencing. **(B)** Bioinformatic pipeline of scRNASeq data treatment. Each step is shown (left column) together with the tool used (right column). **(C)** Number of expressed genes plotted against the number of uniquely mapped reads. (R^2^=0.79 all samples, R^2^= 0.57, selected samples). Black symbols represent cells that passed quality control and grey symbols cells which failed quality control. Circles represent pyramidal neurons under CTRL conditions, squares GABAergic neurons under CTRL conditions. GABAergic neurons in purified neuron cultures are shown in red and in purified neuron cultures treated with OCM in green. **(D)** Mapping statistics of reads for each sample (top, quality control passed, bottom, quality control failed). **(E)** Distribution of reads of transcripts with different origins for each sample (top, quality control passed, bottom, quality control failed).

**Sup. Figure 3:** Comparison of distributions for genes expressed (representative example, sample 34, PYR) within noncoding regions (intergenic, black) and coding regions. Genes were not detected when their expression fell within 98% of intergenic regions. In this case, *Scn1b* and *Kcnc1* were considered to be expressed while *Bsn* was not.

**Sup. Figure 4:** Expression of genes in a cluster with similar expression pattern in different culture conditions. Gene expression was normalized to that of pyramidal cells in control conditions. The number of genes for each cluster is indicated. The continuous black line shows the mean gene expression in different culture conditions.
